# Early detection of white matter hyperintensities using SHIVA-WMH detector

**DOI:** 10.1101/2023.02.03.526961

**Authors:** Ami Tsuchida, Philippe Boutinaud, Violaine Verrecchia, Christophe Tzourio, Stéphanie Debette, Marc Joliot

## Abstract

White matter hyperintensities (WMH) are well-established markers of cerebral small vessel disease (cSVD), and are associated with an increased risk of stroke, dementia, and mortality. Although their prevalence increases with age, small and punctate WMHs have been reported with surprisingly high frequency even in young, neurologically asymptomatic adults. However, most automated methods to segment WMH published to date are not optimized for detecting small and sparse WMH. Here we present the SHIVA-WMH tool, a deep-learning (DL)-based automatic WMH segmentation tool that has been trained with manual segmentations of WMH in a wide range of WMH severity. We show that it is able to detect WMH with high efficiency in subjects with only small punctate WMH as well as in subjects with large WMHs (i.e. with confluency) in evaluation datasets from three distinct databases: MRi-Share consisting of young university students, MICCAI 2017 WMH challenge dataset consisting of older patients from memory clinics, and UK Biobank with community-dwelling middle-aged and older adults. Across these three cohorts with a wide-ranging WMH load, our tool achieved voxel-level and individual lesion cluster-level Dice scores of 0.66 and 0.71, respectively, which were higher than for three reference tools tested: the lesion prediction algorithm implemented in the lesion segmentation toolbox (LST-LPA: Schmidt, 2017), PGS tool, a DL-based algorithm and the current winner of the MICCAI 2017 WMH challenge (Park et al, 2021), and HyperMapper tool (HPM: Mojiri Forooshani et al., 2022), another DL-based method with high reported performance in subjects with mild WMH burden. Our tool is publicly and openly available to the research community to facilitate investigations of WMH across a wide range of severity in other cohorts, and to contribute to our understanding of the emergence and progression of WMH.

**Highlights:**

- We propose a novel 3D Unet-based model, SHIVA-WMH detector, with much improved detection of small WMH across subjects with a wide range of WMH burden compared to existing methods
- We characterize microstructural properties of small white matter hyperintensities in young adults from MRi-Share study

## 1. Introduction

Cerebral small vessel disease (cSVD) represents a spectrum of pathological processes affecting small vessels of the brain. It is a leading vascular cause of dementia and accounts for up to 25% of strokes (Cannistraro et al., 2019; Wardlaw et al., 2013). Most often, cSVD is covert and can be detected on brain imaging of individuals without clinical manifestation of stroke. White matter hyperintensity (WMH) is one of the most well-established imaging markers of cSVD, and is characterized by heightened signal intensity on T2-weighted-fluid-attenuated inversion recovery (FLAIR) sequences of magnetic resonance imaging (MRI). WMH of presumed vascular origin is highly prevalent in neurologically asymptomatic older individuals, and is associated with an increased risk of stroke, cognitive decline, dementia, and mortality (Debette and Markus, 2010; Debette et al., 2019; Wardlaw et al., 2021). So far, the exact etiology and pathogenesis of these age-related WMHs remain elusive, though they are known to be associated with common cardiovascular risk factors, including smoking and hypertension (Moroni et al., 2018). Regardless, they represent an important pre-clinical biomarker of cSVD that should trigger preventive interventions to reduce the risk of stroke and dementia and can be used as a surrogate endpoint in clinical trials (Wardlaw et al., 2021).

Although their prevalence increases with age, small and punctate WMHs have been reported with surprisingly high frequency even in young adults under 40 years of age (Keřkovský et al., 2019; Wadhwa et al., 2019; Wang et al., 2019; Williamson et al., 2018). If they represent early forms of covert cSVD, it is crucial to investigate their emergence and progression in order to study their pathophysiological correlates as well as their association with genetic, environmental, and behavioral risk factors. Clinically, the presence and severity of WMH on MRI are most commonly assessed with visual rating scales, such as the Fazekas (Fazekas et al., 1987) or more anatomically detailed Scheltens Scale (Scheltens et al., 1993) and the Age-Related White Matter Changes (ARWMC) scale (Wahlund et al., 2001). They are all point scales expressing the severity of WMH, with grade 0 indicating no lesion and increasing grade indicating the size and/or number of the lesion, as well as the degree of confluency. While they can be effective for differentiating the most severe cases of WMH from milder cases, these clinical scales are inadequate for the characterization of subjects with the early stages of cSVD, who would at most receive a grade of 1. They also provide very limited information on the spatial extent and distribution of WMH, only distinguishing between WMH found in the periventricular region from deep white matter (for Fazekas) or different lobes in each hemisphere (for ARWMC). It is therefore essential to have automated methods to segment WMH in order to quantify and provide precise spatial information of any lesions in cohorts across the disease spectrum.

While numerous WMH segmentation tools and algorithms exist, including an increasing number of deep-learning (DL) based methods in recent years, there is a lack of automated methods that have been validated in populations with low prevalence and small overall lesion load, hampering the detailed characterization of WMH in young subjects. Indeed, most methods published to date are optimized for detection in older subjects (Guerrero et al., 2018; Li et al., 2018, 2022; Sundaresan et al., 2021; Umapathy et al., 2021) or patients with multiple sclerosis (MS; reviewed in Zeng et al., 2020) who typically manifest a higher load of WMH with large confluent lesions. In these populations, the advantage of more advanced DL-based methods over more traditional signal-processing and machine-learning-based methods is reported to be minimal (Balakrishnan et al., 2021). The true advantage of these advanced methods may be more evident for the segmentation of WMH in subjects with relatively mild lesion load (< 5mL), as recently suggested (Khademi et al., 2021; Li et al., 2022; Rachmadi et al., 2018). However, even those studies evaluating their method in subjects with mild WMH burden with DL-based (Khademi et al., 2021; Rachmadi et al., 2018) or other approaches (Ong et al., 2022; Rachmadi et al., 2020) primarily use databases with 2D FLAIR acquisition with the slice thickness ranging from 3 to 5 mm, which precludes detection of small WMH not on the plane of acquisition. To our knowledge, no study has explicitly optimized the segmentation performance in high-quality 3D FLAIR scans from healthy young-to middle-aged adults with very mild WMH burden.

In the present study, we took advantage of the unique, large neuroimaging database of French university students called MRi-Share (Tsuchida et al., 2021). We first characterize small WMH visible on the 1mm isotropic 3D FLAIR scans in a subsample of 50 MRi-Share participants. Using the multi-shell diffusion-weighted imaging (DWI) also available in these subjects, we demonstrate that albeit their small size and relative sparsity, these small WMH found in young adults already exhibit altered white matter microstructural properties relative to normal-appearing white matter (NAWM). We then describe the development of a 3D Unet-(Ronneberger et al., 2015) based tool, trained with the manually delineated WMH in these subjects. It is based on the model we described previously (Boutinaud et al., 2021) that detected perivascular spaces (PVS), another marker of covert cSVD (Wardlaw et al., 2013; Yu et al., 2022), in the same young subjects. Since we developed this tool in the context of the SHIVA project (https://rhu-shiva.com/), whose aim is to prevent cognitive decline and dementia through a better understanding of cSVD, we call our tool ‘SHIVA-WMH’ detector. Our objective was to create a ready-to-use tool that can chart the emergence and progression of WMH across the adult lifespan in multi-cohort studies with varying age ranges. Thus, in order to cover the full range of age-related WMH severity when tuning the performance of the SHIVA-WMH detector, we also used a publicly available MRI dataset from the MICCAI 2017 WMH segmentation challenge (MWC: Kuijf et al., 2019). We used the openly available 60 training data of the challenge, which were acquired from memory clinic patients in three different institutes and covered a much wider range of WMH severity than the MRi-Share subjects with limited amounts of WMH. We evaluate the performance of our model in the held-out evaluation set of 10 subjects each from the two cohorts. In addition, we manually traced WMH in a small sample of 11 subjects from the UK Biobank data (Alfaro-Almagro et al., 2018), representing research quality scans of community-dwelling middle-aged and older subjects, to be used as the evaluation set coming from an unseen cohort. We compare the performance of the SHIVA-WMH detector with three reference WMH segmentation tools that could be applied out-of-the-box (i.e. without re-training or fine-tuning): 1) LST-LPA (Lesion Prediction Algorithm implemented in the Lesion Segmentation Toolbox; Schmidt, 2017a), a clinical reference tool based on the conventional signal-processing, 2) PGS (Park et al., 2021), a state-of-the-art, ensemble 2D Unet-based model that is currently the winner of the MWC, and 3) HPM (HyperMapper; Mojiri Forooshani et al., 2022), another state-of-the-art 3D Unet-based tool with promising results in subjects with mild WMH burden. We present the performance metrics both at the voxel- and individual lesion cluster-level for each tool, and provide cohort by cohort analysis for performance comparison. Our SHIVA-WMH detector tool with pre-trained models is made openly available at (https://github.com/pboutinaud/SHIVA_WMH) to other researchers to encourage replication and further research into the earliest forms of WMH.

## 2. Material and Methods

### 2.1. Participants and MRI Data description

Table 1 summarizes the key acquisition parameters for the T1-weighted (T1w) and FLAIR images, sample sizes for manually traced WMH, and the range of total lesion load for the three datasets used in the present work, acquired across five different scanners.

**Table 1.** Summary of key acquisition parameters and the range of manually-traced WMH lesion load in the three cohorts. Abbreviations; MWC: MICCAI 2017 WMH segmentation challenge dataset, UMC: the University Medical Center, VU: Vrije Universiteit University Medical Centre, NUHS: the National University Health System, UKB: UK Biobank, TR: repetition time, TE: echo time, TI: inversion time.

| Dataset | Institute | Scanner | T1-weighted voxel size (TR/TE/TI) | FLAIR voxel size (TR/TE/TI) | Train | Test | WMH load range in mL |
| --- | --- | --- | --- | --- | --- | --- | --- |
| MRi-Share | Bordeaux University | 3T Siemens Prisma | 1.00x1.00x1.00mm<sup>3</sup> (2000/2.0/880ms) | 1.00x1.00x1.00mm<sup>3</sup> (5000/394/1800ms) | 40 | 10 | 0 - 1.8 |
| MWC | UMC Utrecht | 3T Philips Achieva | 1.00x1.00x1.00mm<sup>3</sup> (7.9/4.5/--ms) | 0.96x0.95x3.00mm<sup>3</sup> (11000/125/2800ms) | 27 | 3 | 0.8 - 75.0 |
|  | VU Amsterdam | 3T GE Signa HDxt | 0.94x0.94x1.00mm<sup>3</sup> (7.8/3.0/--ms) | 0.98x0.98x1.20mm<sup>3</sup> (8000/126/2340ms) | 27 | 3 |  |
|  | NUHS Singapore | 3T Siemens TrioTim | 1.00x1.00x1.00mm<sup>3</sup> (2300/1.9/900ms) | 1.00x1.00x3.00mm<sup>3</sup> (9000/82/2500ms) | 26 | 4 |  |
| UKB | UKB Imaging Centre | 3T Siemens Skyra | 1.00x1.00x1.00mm<sup>3</sup> (2000/2.01/800ms) | 1.05x1.00x1.00mm<sup>3</sup> (5000/395/1800ms) | -- | 10 | 0.1 - 22.5 |

#### 2.1.1. MRi-Share

The MRi-Share database is a subcomponent of a larger, prospective cohort study on French university students’ health, called iShare (internet-based Student Health Research enterprise, www.i-share.fr). The detailed study and MRI acquisition protocol have been described in Tsuchida et al., (2021). For the training and evaluation of the SHIVA-WMH detector, we used the MRI data and manually-traced WMH from the same sub-sample of 50 subjects described in Boutinaud et al., (2021), drawn from the total sample of 1,867 MRi-Share subjects (mean age 22.1 years, range 18 to 35, 72% female). We also used the T1w and FLAIR data from additional 360 subjects, selected from the 1,817 subjects without any manual tracing of WMH, to enhance the training dataset, as described in the section *SHIVA-WMH detector: training and enhancement*.

All participants were imaged between 2015 and 2018 on a 3T Siemens Prisma MRI scanner (Siemens Healthcare, Erlangen, Germany) with a 64-channel head coil at Bordeaux University, in a single MRI session lasting for ~45 min. It included a 3D T1w magnetization-prepared rapid gradient-echo (MPRAGE) as well as a 3D SPACE FLAIR sequence, both with 1.0 mm isotropic resolution (Table 1). Diffusion-weighted MRI (DWI) data were acquired using a multi-shell multiband x3 sequence with 100 non-collinear diffusion gradient directions (b0/d8 each for anterior-to-posterior (AP) and posterior-to-anterior (PA) phase encoding; b300/d8; b1000/d32; b2000/d60) with the following parameters: TR/TE = 3,540/75 msec; FOV = 118 × 118 mm^2^; 84 slices; 1.7 mm isotropic resolution.

#### 2.1.2. MICCAI 2017 WMH challenge dataset (MWC)

We supplemented the training and evaluation datasets with the MRI and manual tracing of WMH from 60 subjects provided as training datasets in the MICCAI 2017 WMH segmentation challenge (http://wmh.isi.uu.nl/; Kuijf et al., 2019). The set of 3D T1w and 2D or 3D multi-slice FLAIR images were acquired at three different institutes: the University Medical Center (UMC) Utrecht, Vrije Universiteit University Medical Centre (VU) Amsterdam, and the National University Health System (NUHS) in Singapore (20 subjects per site; Table 1). The 3D FLAIR images had been resampled into the transversal direction with a slice thickness of 3 mm by the MWC organizers to make them similar to other 2D FLAIR images in the dataset and to save time for manual annotation. All subjects were recruited at the memory clinics on each site as part of larger cohort studies (UMC Utrecht and VU Amsterdam data from a cohort of 861 subjects, mean age 67.7 years and 46.3% female (Boomsma et al., 2017); NUHS Singapore data from a cohort of 238 subjects, mean age 72.5 years, range 50 to 95 years, 51% female (van Veluw et al., 2015)), and were selected randomly by the MWC organizers.

#### 2.1.3. UK Biobank (UKB)

UK Biobank is the largest cohort study with brain MRI measurements from approximately 50,000 middle-aged and older adults (as of 2022), recruited from communities across the United Kingdom (Miller et al., 2016). All brain imaging data were acquired at one of the three dedicated imaging centers equipped with identical 3T Siemens Skyra scanners (Siemens Healthcare, Erlangen, Germany) with the standard Siemens 32-channel head coil (Alfaro-Almagro et al., 2018). For the present work, we use the raw T1w and FLAIR images acquired using similar sequences as those for MRi-Share (Table 1).

### 2.2. Manual tracing of WMH

#### 2.2.1. MRi-Share

Subjects for manual tracing of WMH were selected based on the visual inspection of a neuroradiologist (BM) who reviewed the raw T1w and FLAIR images of the entire dataset, to cover varying degrees of both WMH and visible PVS, from no detectable WMH or PVS to many visible WMH (>10) and/or PVS. A trained investigator (AT) then performed voxelwise manual segmentation of each WMH on the raw FLAIR images using Medical Image Processing, Analysis and Visualization (MIPAV) software (v 7.4.0).

Specifically, WMH was segmented on each axial slice of the FLAIR image, viewed along with coronal and sagittal views using the 3D view setting of the MIPAV to check the 3D shape and extent of hyperintense signals. Any punctate region of increased intensity within the white matter, as well as hyperintense rims around the ventricles that were thicker than 2 mm, were segmented. This included punctate hyperintense regions sometimes found around the PVS visible on T1w images. For each hyperintense region found, the ‘paint grow’ tool of the MIPAV was applied to automatically paint every neighboring voxels that have a higher intensity than and within a 3 mm distance from the selected voxel. Following the initial segmentation of the first ten subjects, they were reviewed and modified by a second expert (LL). Any discordance between the two raters was then reviewed together to reach a consensus. Subsequently, the remaining 40 MRI datasets were manually segmented by the first expert only.

#### 2.2.2. MWC

We used the publicly available manual tracing of WMH for the 60 MWC training dataset. The details of the procedure are described in Kuijf et al., (2019). Briefly, manual tracing of WMH, as well as any other pathologies, was performed by two expert raters by consensus, in which the tracing performed by the first rater was reviewed by the second rater and corrected by the first rater.

#### 2.2.3. UKB

A small sample of 11 subjects was selected to cover a range of estimated WMH load (0.7 to 16mL, representing values in the first and last decile of the entire dataset) from a pool of 13554 subjects (available at the time of the selection) with ‘usable’ quality T1w and FLAIR images as well as the WMH load estimated by the Brain Intensity Abnormality Classification Algorithm tool (BIANCA: Griffanti et al., 2016). The same rater who performed the manual tracing of WMH for the MRi-Share database (AT) manually traced WMH on the raw FLAIR images using the 3D Slicer tool (version 4.11.20210226: https://www.slicer.org), using the same criteria applied during the segmentation of WMH for MRi-Share. With the 3D Slicer tool, the ‘Threshold’ effect in the Segment Editor module was used to specify an intensity range that preliminarily isolated visible hyperintensities from the surrounding white matter in any given location. As in MRi-Share, each axial slice was reviewed slice by slice, together with coronal and sagittal views to check the entire 3D extent of each lesion, and the ‘Paint’ tool that painted regions with the specified intensity range was used to segment individual hyperintensities deemed as lesion, in each plane of the 3D view.

### 2.3. Comparison of microstructural properties inside WMH to normal-appearing white matter (NAWM) in MRi-Share

The processing pipeline of DWI data to obtain diffusion tensor imaging (DTI: Basser et al., 1994) and neurite orientation dispersion and density imaging (NODDI: Zhang et al., 2012) metrics in MRi-Share has been described in detail in Tsuchida et al., (2021). Briefly, DWI data were preprocessed with the Eddy tool from FMRIB Software Library (FSL, version 5.0.10: https://fsl.fmrib.ox.ac.uk/fsl) to correct for susceptibility and eddy-current distortion, then denoised using the non-local means filter (Coupe et al., 2011, 2008) using *nlmeans* tool implemented in the *Dipy* package (0.12.0: Garyfallidis et al., 2014). The DTI model was fit using the *Dipy* package, while the NODDI model was fit using the *AMICO* tool (Daducci et al., 2015).

The resulting scalar images of DTI and NODDI metrics were coregistered to the native T1w image space using *antsRegistrationSyNQuick* script of Advanced Normalization Tools (ANTs, version 2.1: http://stnava.github.io/ANTs/) package. For the present work, we focused on the neurite density index (NDI) from NODDI, fractional anisotropy (FA), and mean diffusivity (MD) from DTI metrics, which all have been shown to be altered in WMH in MS (Alotaibi et al., 2021) or in older subjects (Muñoz Maniega et al., 2015; Riphagen et al., 2018).

To compare NDI, FA, and MD values inside WMH and NAWM in the 50 MRi-Share subjects with the manual tracing of WMH, manually traced WMH masks in the native FLAIR space were coregistered linearly to the T1w image by applying the transformation matrix generated by the coregistration of FLAIR to T1w with *flirt* tool from the FSL (Jenkinson et al., 2002). The mask of NAWM was generated by combining both the manually traced WMH and PVS masks, then subtracting this from the cerebral white matter labels generated by the Freesurfer (v6.0: http://surfer.nmr.mgh.harvard.edu/) in the native T1w space. To remove any partial volume effects near the border of lesion or other tissue types (grey matter and cerebrospinal fluid), the mask of cerebral white matter was eroded once using *fslmaths* tool from the FSL. To avoid the partial volume effects of the cerebrospinal fluid in the WMH mask, periventricular lesion clusters within a 2 mm distance of individual ventricle maps (generated with Freesurfer v6.0) were removed from the analysis. Forty-six out of 50 subjects had non-empty WMH masks and could be included in the analysis. Mean values of NDI, FA, and MD inside the resulting WMH and NAWM masks were computed and compared by performing a within-subject *t*-test for each metric.

All paired *t*-tests were performed in *R*, version 4.2.2 (R Core Team, 2018), and visualized using *ggpubr* package (Kassambara, 2022).

### 2.4. SHIVA-WMH detector

#### 2.4.1. Preprocessing

In order to prepare the input image arrays for the SHIVA-WMH detector, the following preprocessing steps were performed on the T1w, FLAIR, and manually-traced WMH masks:

1. Reorientation to match either LAS or RAS (left or right/anterior/superior) orientation using *fslreorient2std* tool from the FSL.
2. For the MWC dataset, images were resampled to 1mm isotropic using *flirt* from the FSL (Jenkinson et al., 2002), with -applyisoxfm and -noresampleblur options.
3. Linear coregistration of FLAIR to T1w image was performed with *flirt* (Jenkinson et al., 2002) for MRi-Share and UKB datasets, and the generated transformation matrices were applied to the WMH masks to bring them to the reference T1w images. This step was skipped for the MWC dataset, which had already been aligned by the organizers.
4. A brain mask created based on the individual T1w image was used to obtain a bounding box around the brain (centered on the mass center of the brain mask), and to crop all images to a uniform dimension of (160 x 216 x 176 voxels).
5. Voxel intensity values inside the brain mask were linearly rescaled to values between 0 and 1 by setting the 99th percentile value as the maximum and setting any higher intensity values as 1.

#### 2.4.2. SHIVA-WMH architecture and implementation

Our model is based on the previously published PVS detector (Boutinaud et al., 2021) with an Unet-like architecture of Ronneberger et al., (2015). We made the following modifications to improve performance or to adapt it for the specific task of WMH segmentation;

- For the primary multi-channel model, the input layer was modified to accept multi-modal input of T1w and FLAIR images.
- Pretraining with auto-encoder was not performed since the larger training set with multi-modal input in the present work allowed relatively fast training without the pretraining with auto-encoder.
- The architecture of the network was modified to have an increased number of initial kernels (feature maps, nK_init_), from 8 to 10, and the multiplication factor (mF) applied to the number of kernels at the first convolution layer of each stage (or depth) after the first was slightly reduced from 2 to 1.8.
- The dropout rate applied at each stage (after the max pooling for encoding or after the last convolution for decoding blocks) was increased from 0.1 to 0.5, except in the first encoding block, which had a reduced dropout rate of 0.05 to increase the rate of retention, reflecting the general recommendation for optimal dropout rate across a wide range of networks and tasks (Srivastava et al., 2014).
- Convolution blocks now use a Swish activation function (*equation (1)*) rather than the rectified linear unit (ReLU; Glorot and Bengio, 2010) used previously, since it has shown an advantage over ReLU on deeper models across a number of challenging datasets (Ramachandran et al., 2017).

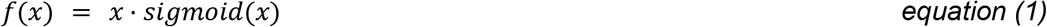

Figure 1 provides a schematic overview of the resulting architecture of the SHIVA-WMH model. The modified model has slightly less trainable parameters (40 million rather than 44 million in the Boutinaud et al. study) and also extracts a higher number of features at higher resolution upper stages and less at deeper levels, which we found to be advantageous for the detection of small lesions like PVS and small WMH.

**Figure 1.**
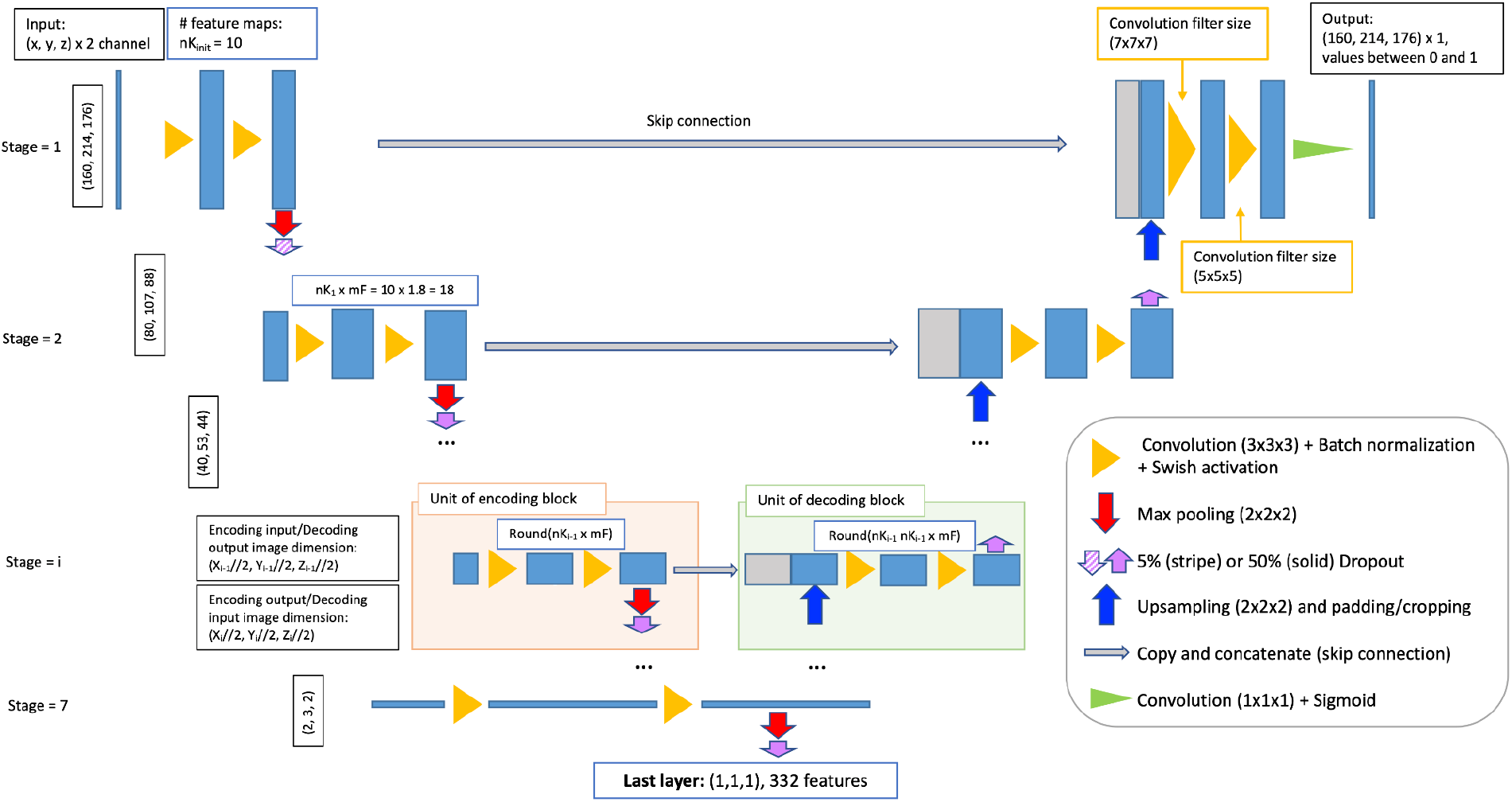
Schematic overview of the SHIVA-WMH detector network architecture. The figure illustrates the 3D Unet architecture used for the SHIVA-WMH detector. Each blue box corresponds to the multi-channel feature maps, with the number of features at each stage indicated in the white box with blue outline. The gray boxes are copied and concatenated feature maps from the encoding path to the decoding path. The arrows and triangles stand for different operations as indicated inside right legend.

In addition to the network architecture modifications, we performed the following data augmentations to the training dataset to increase the model robustness: (i) flipping on the midsagittal plane, (ii) voxel translations (up to plus or minus five voxels in X, Y, and X axes), (iii) non-linear voxel intensity value transformation using a Bézier curve, in the similar fashion as in (Zhou et al., 2021), with two endpoints set to [0, 0] and [1, 1] and two control points within this range generated randomly. The probability of each type of augmentation for an image at a training epoch was set to 0.5, 0.9, and 0.9, respectively. In particular, the non-linear voxel intensity value transformation was found to significantly improve the generalizability of the detector when predicting lesions in unseen datasets (unpublished observation from the PVS detector).

We implemented the network in Python 3.7, using *Tensorflow* 2.7 with *Keras* backend, *scikit-learn* (1.0.1), and *scikit-image* (0.18.3). The network was trained on a computer (Ubuntu 22.04) with a Xeon ES2640, 40 cores, 256 Gb RAM, and a Tesla V100 GPU with 32 Gb RAM; inferences can be done on any GPU compatible with the Tensorflow version with at least 8Gb RAM. We used a 5-fold cross-validation scheme to train the network, stratified on WMH voxel load and cohort. We used the Adam optimizer with the default parameters of *beta1* = 0.9, *beta2* = 0.999, *epsilon* = 1e–7, and used a cyclical learning rate with exponential decay, with the initial and maximum learning rates set to 1e-6 and 0.001, respectively. As in the Boutinaud et al. study, we used a Dice loss function (*equation 2*) as a loss function:

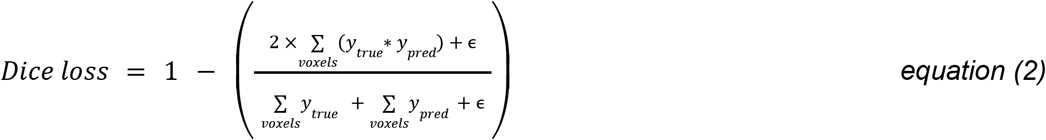

where *y_true_* and *y_pred_* represent the image arrays for the ground truth and predicted WMH, respectively, and (*y_true_* * *y_pred_*) is an element-wise multiplication of the two arrays, representing the intersection between the two images. We used the smoothing constant *ϵ* of 1e-6 to prevent the division by 0. We used a batch size of 4 for each training fold to fit in the available GPU memory. The output maps from each fold, valued between 0 and 1, were averaged to create the final WMH prediction map, also valued between 0 and 1.

#### 2.4.3. Training and enhancement

We initially trained our model with T1w and FLAIR images from 40 MRi-Share and 50 MWC subjects (Table 1), using the manually-traced WMH. We call this model our “base” model. However, due to the large imbalance in the total number of voxels labeled as WMH in the two cohorts (only ~8K voxels of WMH collectively in MRi-Share compared to ~777K voxels in MWC training data; also see Figure 2) the Dice loss function would inevitably bias the optimization towards models that can detect larger WMH lesions. In order to gauge how much the presence of MWC training data degrades the performance of our model in MRi-Share subjects, we trained another model with only MRi-Share training data, which we call “MRi-Share-specific” model. As shown in the Supplemental Figure 1, the comparison of segmentation accuracy in the MRi-Share subjects from the validation sets in each training fold indicated lower accuracy of the “base” model relative to “MRi-Share-specific” model, as expected.

**Figure 2.**
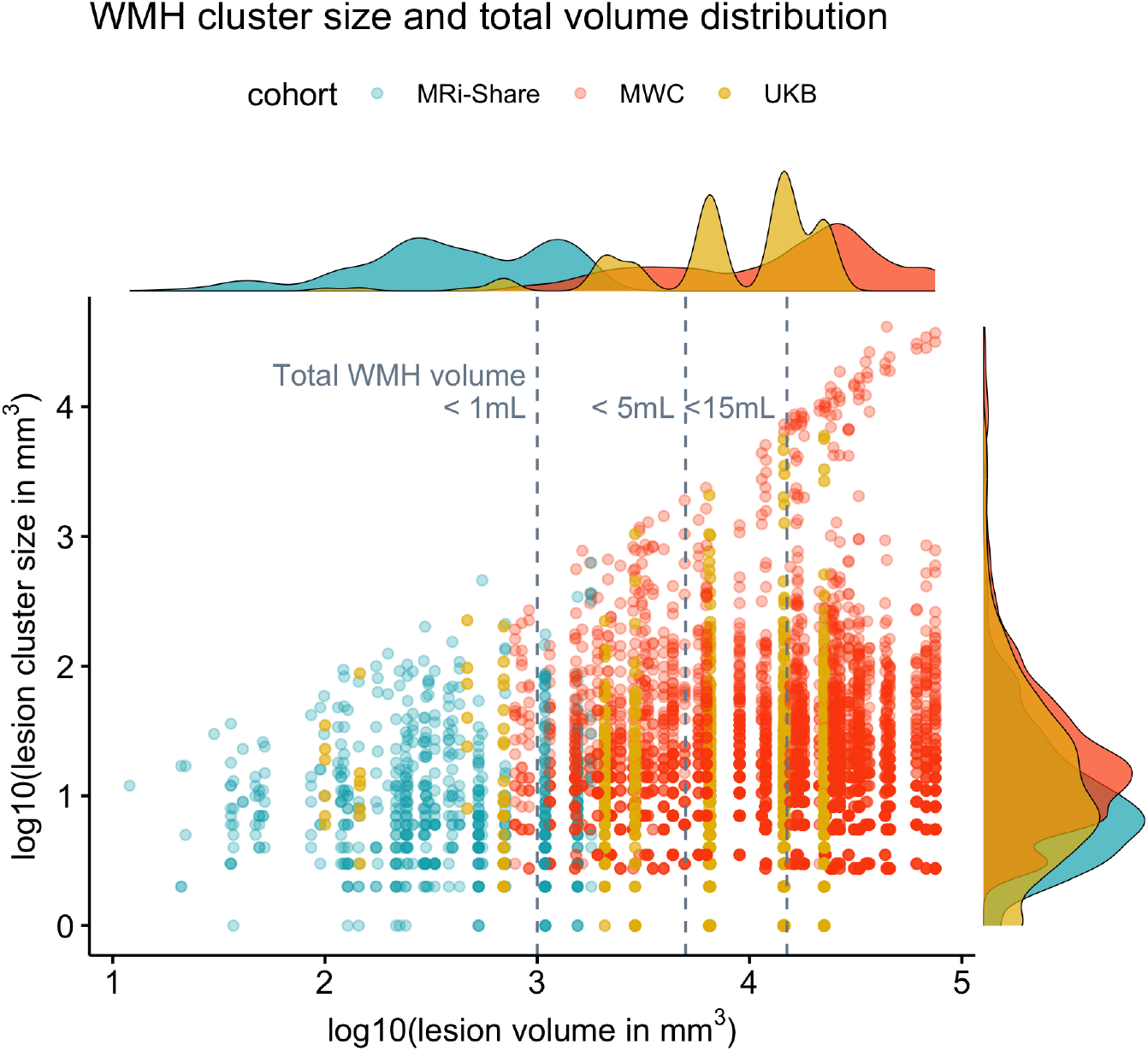
Distribution of total lesion load and individual lesion size of the manually delineated WMH in the three datasets. The x-axis plots the log10-transformed total WMH volume (in mm^3^) against the y-axis showing the distribution of the log10-transformed individual lesion cluster size (in mm^3^) in each subject, based on the manually-traced WMH (i.e each dot along the given x-value representing the lesion clusters in a single subject). Colors indicate subjects from each cohort (MRi-Share in *turquoise*, MWC in *orange*, and UKB in *gold* colors). Histograms on the margins show the distribution of the total lesion volume (top) or the lesion cluster size (right) separately for each cohort.

To overcome the problem of imbalances, we took the advantage of the large pool of still unannotated MRi-Share dataset to perform semi-supervised learning approach using the predictions of “MRi-Share-specific” model that could segment small WMH with relatively high accuracy (see 4.2.2 for a review of such approach: Tajbakhsh et al., 2020). We first obtained the predictions of WMH in 100 randomly selected MRi-Share subjects out of 1,817 subjects without the manual tracing of WMH. We filtered out those with very low or high predicted load of WMH by removing subjects in the top and bottom 5 percentile of the total estimated WMH volume, and used the WMH labels in the 90 subjects to supplement the original training data from 90 subjects (“enh90” model, for enhancement with 90 predicted data).

While the “base” model used the Glorot uniform initializer to initialize its weights, “enh90” model used the weights of the “base” model as the initial weights to speed up the training. Since it showed some indication of improvement over the “base” model in the validation sets of both MRi-Share and MWC training data (Supplemental Figure 1), we repeated the process with an additional enhancement using a new set WMH labels from 270 MRi-Share subjects (“enh270” model), using the weights of “enh90” model as the initial weights. This model showed further improvement in the MRi-Share validation sets and similar performance in the MWC. We repeated the process with increasing numbers of enhancements (300 and 400 additional training labels), each using the weights of the preceding model as the initial weights. However, since there were no signs of further improvement in the MRi-Share validation sets after the “enh270” model (Supplemental Figure 1), we selected this model as the optimal detector; it will be referred to in the following as the SHIVA-WMH detector.

#### 2.4.4. FLAIR-only version

We initially focused on developing a model that uses both T1w and FLAIR as inputs, following an earlier observation of the superior performance of multi-modal over FLAIR-only models. However, to formally compare the impact of having only FLAIR as an input and to potentially allow quantification of WMH in datasets with only FLAIR images, we created the modification of the SHIVA-WMH detector with FLAIR only input. The training process of the FLAIR-only version and its performance comparison with the multi-modal version is described in the Supplemental Material.

### 2.5. Performance evaluation

We evaluated the performance of the SHIVA-WMH detector and existing methods in the held-out test dataset from the three cohorts, including ten unseen data each from MRi-Share and MWC and 11 data from UKB. The UKB test data represent a dataset coming outside of cohorts used in training (Table 1), with the levels of WMH severity being the intermediate between those of MRi-Share and MWC.

#### 2.5.1. Evaluation metrics

We focused on the metrics that quantify the spatial similarity of the ground truth and predicted WMH both at the voxel- or individual lesion cluster-level. Specifically, we counted the number of true positives (TP), false negatives (FN), and false positives (FP) voxel-by-voxel or at the level of individual lesion clusters (each individual lesion cluster defined as a 3D connected component using voxel connectivity of 26) to compute the following performance metrics.

*Voxel-level or cluster-level true positive rate (VL- or CL-TPR)*: It is measured as the number of TP voxels or lesion clusters divided by the number of ground truth voxels or clusters (i.e. TP + FN), and is equivalent to *sensitivity* or *recall*.
*Voxel-level or cluster-level positive predictive value (VL- or CL-PPV)*: It is measured as the number of TP voxels or clusters divided by the number of predicted WMH voxels or clusters (i.e. TP + FP), and is equivalent to *precision*.
*Voxel-level or cluster-level Dice coefficient (VL- or CL-Dice)*: It is the harmonic mean of the *TPR* and the *PPV*, or, equivalently, it can be expressed as;

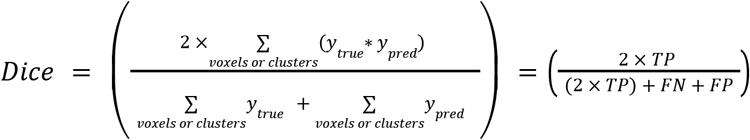

Note that the term *F1* score is used sometimes to refer to lesion cluster-based metric (*CL-Dice* in our terminology) to distinguish from the *VL-Dice*, which is often referred as *Dice score* or *Dice similarity coefficient (eg. Kuijf et al., 2019)*, even though mathematically the Dice and F1 scores are equivalent (Reinke et al., 2021).

We also report the modified Hausdorff distance (95th percentile: HD95) as a measure of the accuracy of the segmentation boundaries. It is defined as the 95th percentile of the symmetric surface distances (Hausdorff distance) of two binary images, with lower values indicating shorter overall distances between the two images. We used the implementation of HD95 computation from the MedPy package (version 0.4.0).

#### 2.5.2. Metric comparisons with existing methods

We performed sets of paired *t-tests* with subject as the within-factor that compared each performance metric of the SHIVA-WMH against the LST-LPA, PGS, and HPM, separately for each of the three cohorts in the held-out test set to allow evaluation of performance in the cohorts with very different demographic and WMH lesion characteristics. Note that PGS performance could not be compared for the MWC test subjects, since they were part of the training data used for this tool. Each comparison was Bonferroni corrected for the number of comparisons made for the given cohort (three for MRi-Share and UKB, two for MWC). All paired *t-*tests were performed in *R*, version 4.2.2 (R Core Team, 2018), and visualized using *ggpubr* package (Kassambara, 2022). Summary tables were generated using *gt* package (Iannone et al., 2020).

For all metric computations, prediction maps were thresholded at 0.5 to make a fair comparison with the PGS tool, whose output prediction maps were already thresholded at this value. For all other tools (SHIVA-WMH, LST-LPA, HPM), lower thresholds improved the VL- and CL-Dice scores slightly (thresholds that resulted in the highest average VL- and CL-Dice scores were 0.2 for SHIVA-WMH, 0.1 for LST-LPA and HPM), but the overall patterns remained essentially the same (not shown).

### 2.6. Comparison algorithms

#### 2.6.1. Lesion Segmentation Toolbox - Lesion Prediction Algorithm (LST-LPA: Schmidt, 2017b)

The LST-LPA is an open-source MATLAB tool that uses only a FLAIR image as an input to segment WMH without requiring any optimizations or re-training. It is based on a conventional signal processing method with binary classification using a logistic regression model. While it was originally developed to detect WMH in MS patients (Schmidt et al., 2012), it has been applied in the context of age-related WMH and widely used (Garnier-Crussard et al., 2020; Ribaldi et al., 2021; Vanderbecq et al., 2020). Being part of the toolbox for the SPM software, it is fully automated and simple to use, and is often selected as a reference tool for the new WMH detection method development (Balakrishnan et al., 2021). Since it comes with a built-in preprocessing pipeline that includes intensity normalization, we used the raw FLAIR images from the test dataset as the input to obtain the predicted maps of WMH.

#### 2.6.2. PGS (Park et al., 2021)

The PGS tool is a 2D Unet-style model with a multi-scale highlight foreground method to augment the influence of small lesions or voxels lying in the lesion boundaries, and is currently the best-ranking method that has been submitted to the MICCAI 2017 WMH challenge. The pre-trained model submitted to the challenge is available as the docker-contained code from the challenge website (https://wmh.isi.uu.nl/results/pgs/). It has been trained on the 2D axial slices of T1w and FLAIR images from the same MWC dataset used in the present study. Because our MWC test dataset is part of the training dataset in the challenge (and therefore in the PGS model), evaluation of this tool in the MWC test dataset was not performed. Since this tool had been trained on the bias-field corrected data prepared by the challenge organizers (using SPM12), we used the bias-field corrected T1w and FLAIR images of the MRi-Share (based on SPM12, as described in Tsuchida et al., (2021)) and UKB datasets (based on FSL FAST, as described in Alfaro-Almagro et al., (2018)) as inputs for this tool.

#### 2.6.3. HyperMapper (HPM: Mojiri Forooshani et al., 2022)

The HPM tool is a 3D Unet model with Monte Carlo dropout layers incorporated in the encoding layers, and reported high accuracy on a large and diverse test dataset representing 160 subjects from seven different studies. Of particular interest from our perspective, it also reported very high accuracy on the subsample of 50 test subjects with mild WMH cases with the average of 2 mL of lesions. Unlike the PGS tool and many other recently published methods (including all other tools submitted to the MICCAI 2017 WMH challenge), the HPM did not use the MWC dataset to train the models. As such, its performance on the MWC test dataset could be evaluated against the SHIVA-WMH. The HPM tool is publicly available from https://github.com/AICONSlab/HyperMapp3r. We used the docker-contained image included in the repository to perform predictions on the test datasets. Since this tool also has been trained on the bias-fileld-corrected training data, we used the bias-field corrected T1w and FLAIR images for all test datasets.

## 3. Results

### 3.1. Characterization of manually-traced WMH in the 3 cohorts

The sample of subjects with manual segmentations of WMH in the present study comprised 50 young adults (18-35 years of age) from the MRi-Share study, 11 middle-aged to older UKB participants (>40 years of age), and 60 memory clinic patients (approximately >50 years of age) from the MWC. Figure 2 shows the distribution of total WMH lesion volume and individual lesion cluster sizes of each participant (with the x-axis effectively ordering every subject according to the total lesion volume) in each of the three cohorts with the manual tracing of the WMH. It demonstrates the vastly different scales of the overall amount of WMH lesions in the young subjects of the MRi-Share from the older, memory clinic patients represented by the MWC: without the log10-transformation of the scales, the total WMH volumes for the MRi-Share subjects would cluster around 0, since most subjects have less than 1 mL of WMH in total. Representing middle-aged and older subjects, the eleven subjects from the UKB show levels of WMH load intermediate between those of the MRi-Share and MWC subjects. Expectedly, the maximum size of individual lesion increases with the total lesion volume, with very large lesions in subjects with the most severe cases of WMH, likely representing the confluence of deep and periventricular WMH. Despite that, it should be noted that subjects with high WMH load also have many relatively small lesion clusters, which underscores the importance of accurately segmenting small lesions for comprehensive characterization of WMH.

### 3.2. Comparison of microstructural properties inside WMH to normal-appearing white matter (NAWM) in young subjects of MRi-Share

Despite their relative sparsity and small sizes, within-subject comparisons of white matter properties inside and outside the manually traced lesions revealed significant differences in NDI, FA, and MD values in the WMH found in MRi-Share participants: compared to NAWM, WMH showed a decreased NDI (mean difference [95% confidence intervals] = −0.179 [−0.145, −0.213], paired *t*-test *p* < 0.0001) and FA (−0.125 [−0.104, −0.144], paired *t*-test *p* < 0.0001) values, and an elevated MD values (+2.2×10^-4^ [1.88×10^-4^, 2.57×10^-4^] mm^2^/sec, paired *t*-test *p* < 0.0001) (Figure 3). The changes in NDI, FA, and MD values were visible at the level of individual lesion clusters in some cases, as in the example shown in Figure 3.

**Figure 3.**
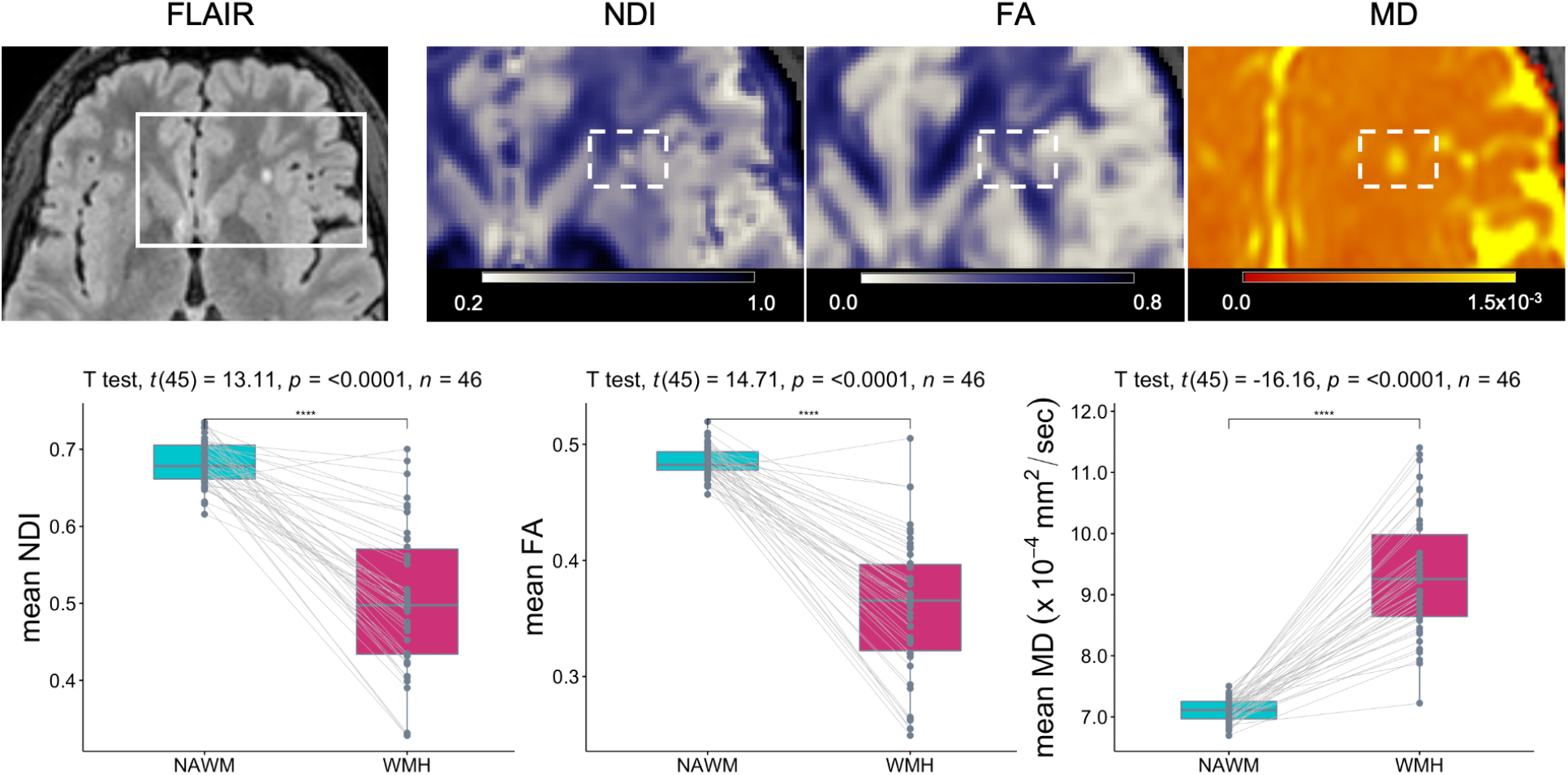
White matter microstructural properties of small WMH in MRi-Share. The top row shows an example axial slice of a FLAIR image of an MRi-Share subject showing an isolated punctate WMH, together with blown-up images of NDI, FA, and MD maps in the same slice to highlight the decreased NDI and FA values and increased MD values within the WMH (in the dotted rectangles). The bottom row shows the within-subject comparisons of mean NDI, FA, or MD values inside the NAWM and WMH in 46 subjects with at least one WMH after removing WMH too close to ventricles to avoid partial volume effects from the cerebrospinal fluid in the ventricles.

### 3.3. Detection of WMH with SHIVA-WMH detector across a wide range of lesion loads

We used the manually traced lesions from both MRi-Share and MWC, and enhanced training data from additional MRi-Share participants to train the SHIVA-WMH detector. We then evaluated the performance of our detector against three reference methods (LST-LPA, PGS, and HPM), in the held-out evaluation test subjects comprised of 10 MRi-Share, 11 UKB, and 10 MWC subjects. Table 2 shows the summary of performance metrics for each method across all 31 test subjects, except for PGS, which was not evaluated for MWC test subjects. Overall, they indicate the superior segmentation accuracy of the SHIVA-WMH detector over the three reference methods (LST-LPA, PGS, and HPM), with higher sensitivity (TPR) and precision (PPV) both at the voxel- and lesion cluster-level than any of the reference methods, resulting in the significanyl higher VL- and CL-Dice scores (all paired t-tests *p* < 0.001 for VL-Dice and *p* < 0.0001 for CL-Dice). The correlation between the log-transformed volume of manually traced lesion and segmented WMH across the test set subjects was also the highest for SHIVA-WMH compared to the three reference methods (Figure 4). Although FLAIR-only version of SHIVA-WMH had slightly worse performance than the primary multi-modal input version, it still had nominally better VL-Dice scores and significantly better CL-Dice scores against all three reference methods tested (*p* < 0.0001 for CL-Dice; Supplemental Table 1).

**Table 2.**
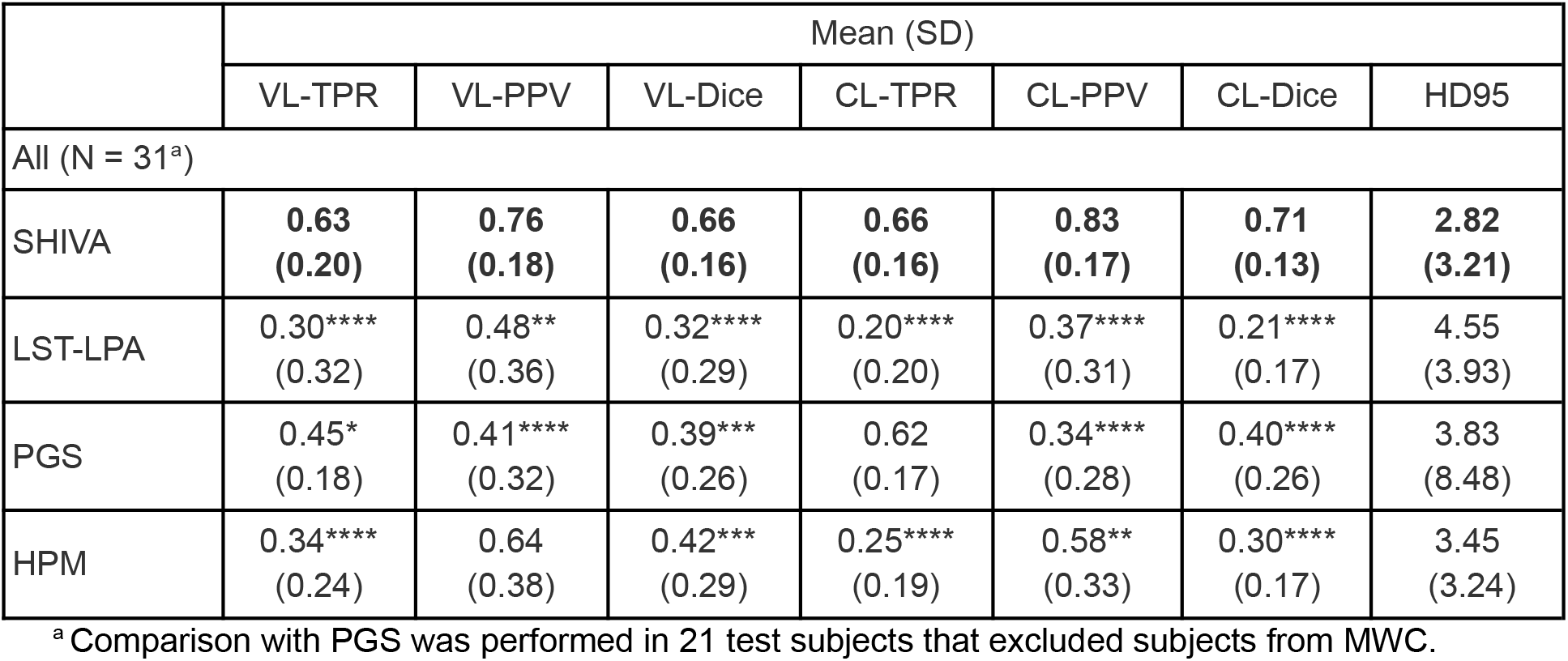
Comparison of SHIVA-WMH against three reference methods across the 31 test subjects for each performance metric. Mean and standard deviations (SD) of each metric across all the test subjects are shown for SHIVA-WMH and the three reference methods (LST-LPA, PGS, HPM). For each metric, best scores are indicated in bold. Asterisk indicates the degree of statistical significance for each paired-t test comparing SHIVA-WMH against each of the reference methods: **** *p* < 0.0001, *** 0.0001 ⩽ *p* <0.001, ** 0.001 ⩽ *p* <0.01, * 0.01 ⩽ *p* <0.05.

**Figure 4.**
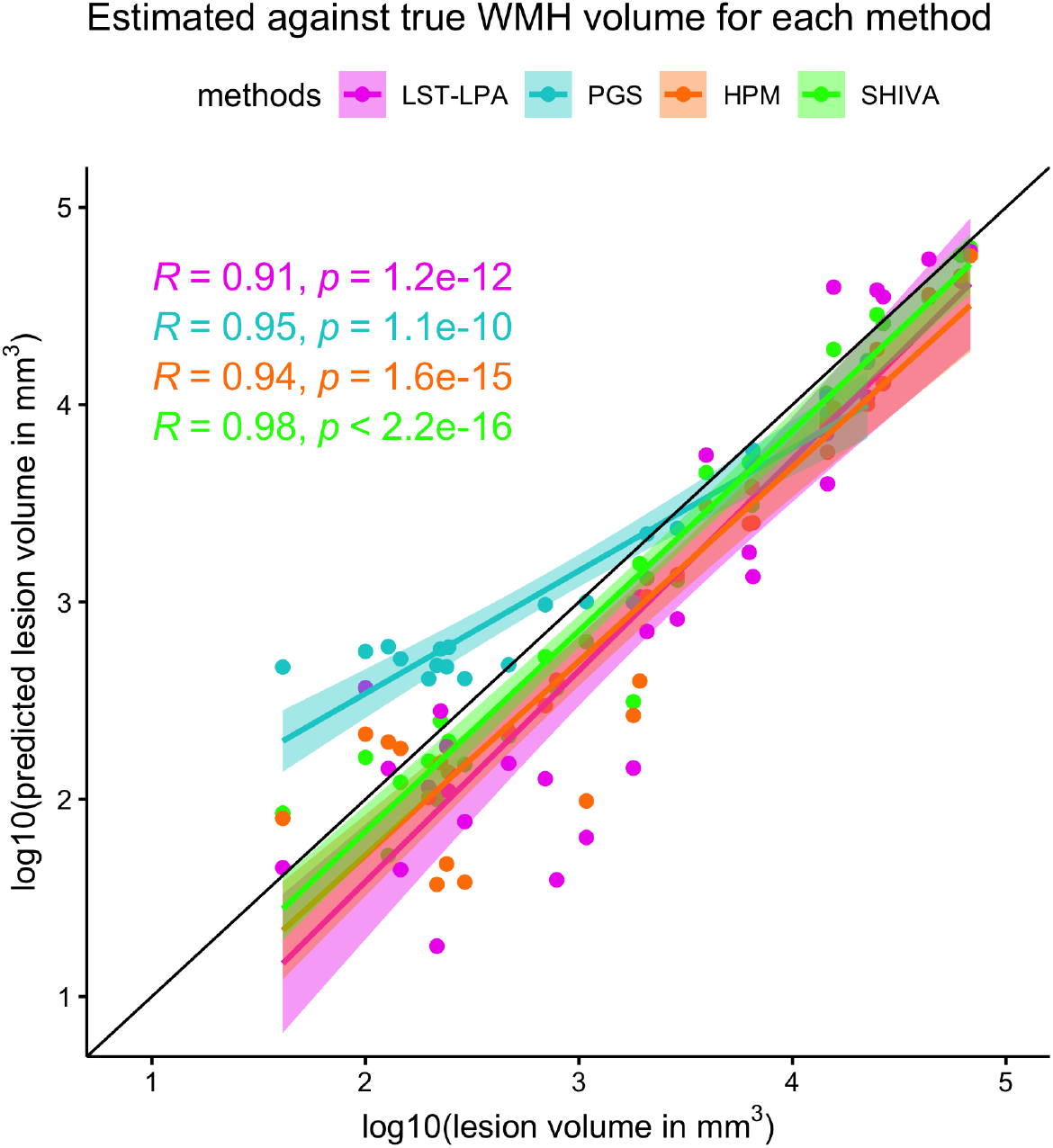
Correlations between total WMH volume of the manually delineated lesions and WMH segmented by each method across the test-set subjects. The x-axis plots the log10-transformed total WMH volume (in mm^3^) based on the manually-traced WMH against the y-axis showing also the log10-transformed total volume (in mm^3^) of WMH segmented by each method. Each dot along the given x-value represents a single subject, with different colors indicating WMH volume estimates based on each method (LST-LPA in *pink*, PGS in *turquoise*, HPM in *orange*, and SHIVA-WMH in *green*). Regression lines with confidence intervals are also shown for each method in respective colors, as well as Pearson’s correlation coefficients (*R*) between the log10-transformed volumes and associated *p* values.

Because the evaluation test set comprised subjects with very different demographics and WMH burden, we performed more detailed analysis by comparing the performance metrics separately for each of the three cohorts. Figure 5 graphically illustrates quantitative comparisons of the two main metrics of interest, VL- and CL-Dice scores, in each cohort. Supplemental Figure 2 shows similar cohort-specific comparisons for FLAIR-only version of SHIVA-WMH. Table 3 provides the same summary as in Table 2, but separately for each cohort. Finally, qualitative comparisons of example segmentations from each method are shown in Figure 6, separately for representative test set subjects with either only small WMH lesions (mild) or with large confluent lesions (severe) from each cohort (except for MRi-Share, in which none of the subjects had severe WMH).

**Figure 5.**
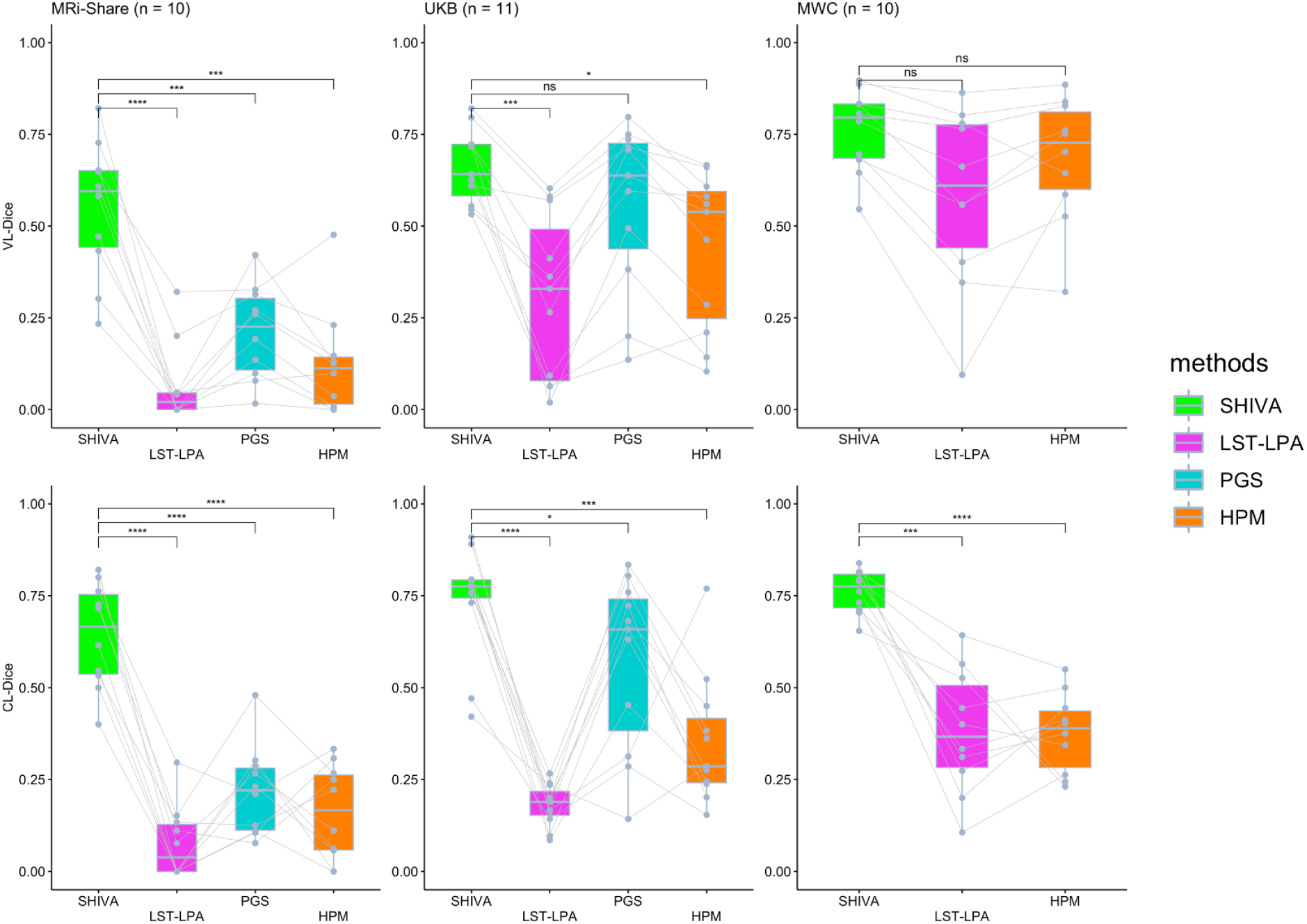
Voxel (VL-) and Cluster-level (CL-) Dice scores of SHIVA-WMH compared with LST-LPA, PGS and HPM tools in the test-set subjects in each cohort. Comparisons of VL-Dice (*top row*) and CL-Dice (*bottom row*) scores between SHIVA-WMH against the reference tools (LST-LPA, PGS, and HPM) are shown, separately for MRi-Share (n = 10), UKB (n = 11), and MWC (n = 10) test subjects. Asterisk indicates the degree of statistical significance for each paired-t test comparing SHIVA-WMH against each of the reference methods: **** *p* < 0.0001, *** 0.0001 ⩽ *p* <0.001, ** 0.001 ⩽ *p* <0.01, * 0.01 ⩽ *p* <0.05, ns *p* ⩾ 0.05.

**Figure 6.**
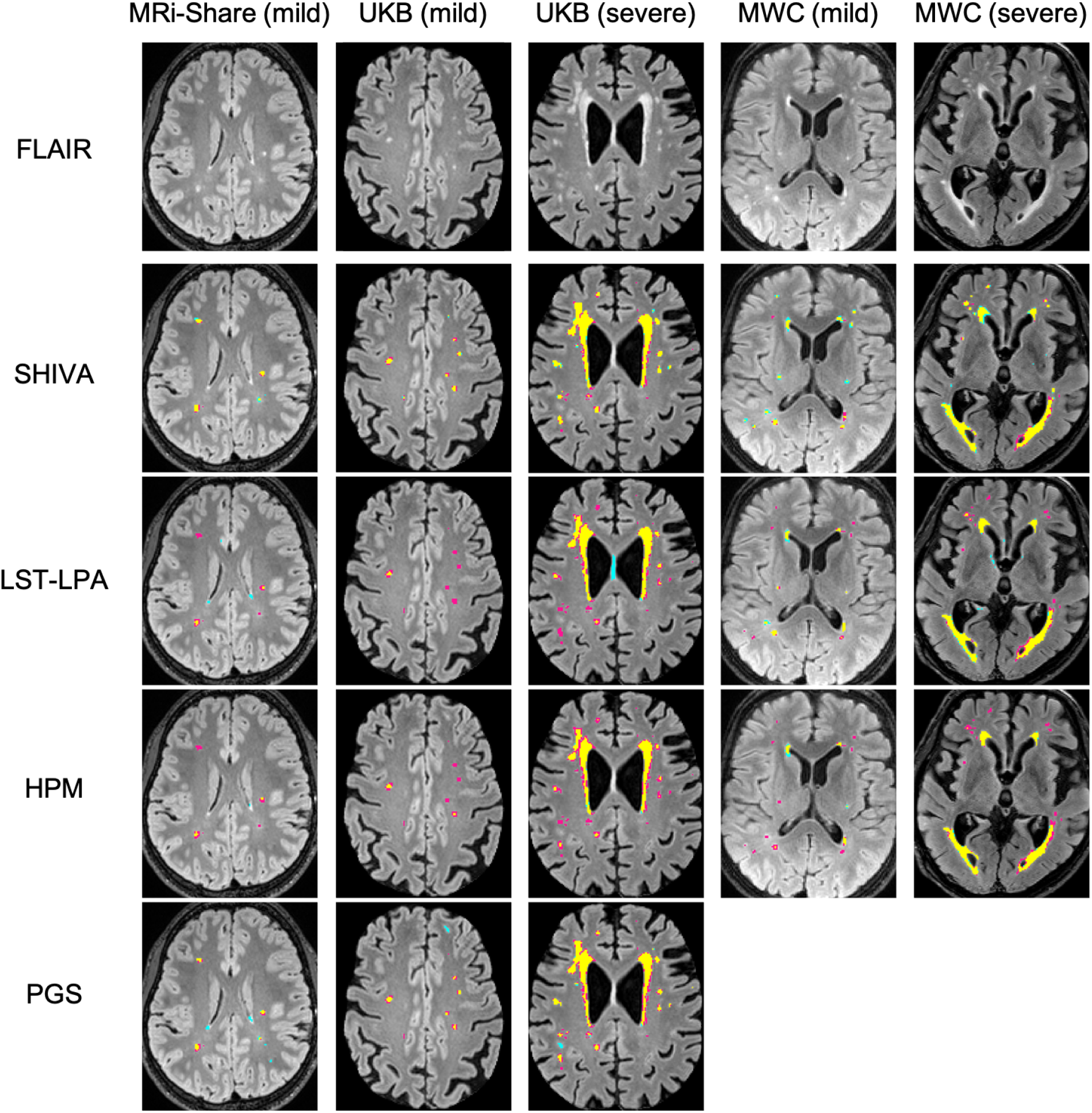
Examples of segmentation output by SHIVA-WMH detector, LST-LPA, PGS, and HPM tools. Examples of WMH segmentations by SHIVA-WMH, LST-LPA, PGS, and HPM are shown separately for representative subject(s) in each cohort with either mild (< 5mL) or severe (> 5mL) WMH load. The top row shows the selected axial slices from each subject, and second to last rows show the segmentation results of each tool, with *yellow*, *pink*, and *cyan* colors indicating true positive, false negative, and false positive voxels, respectively.

**Table 3.**
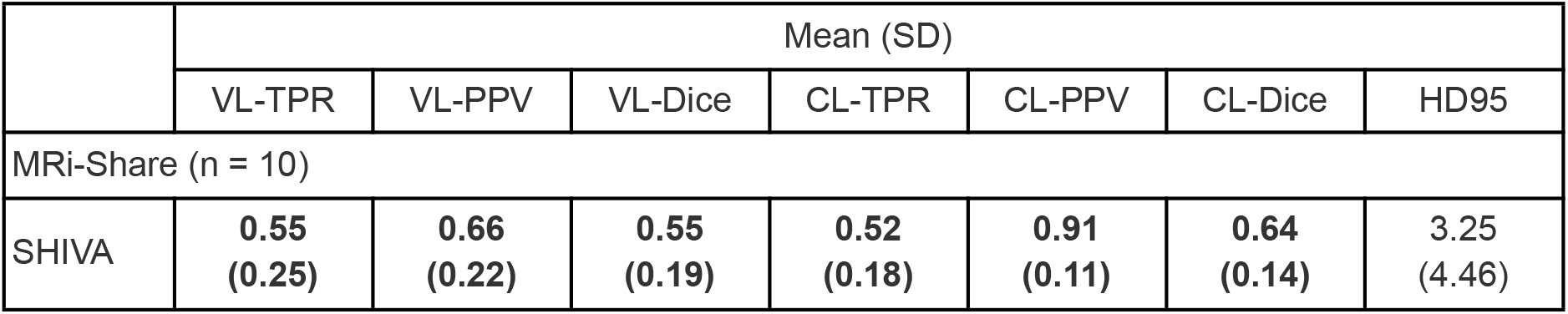
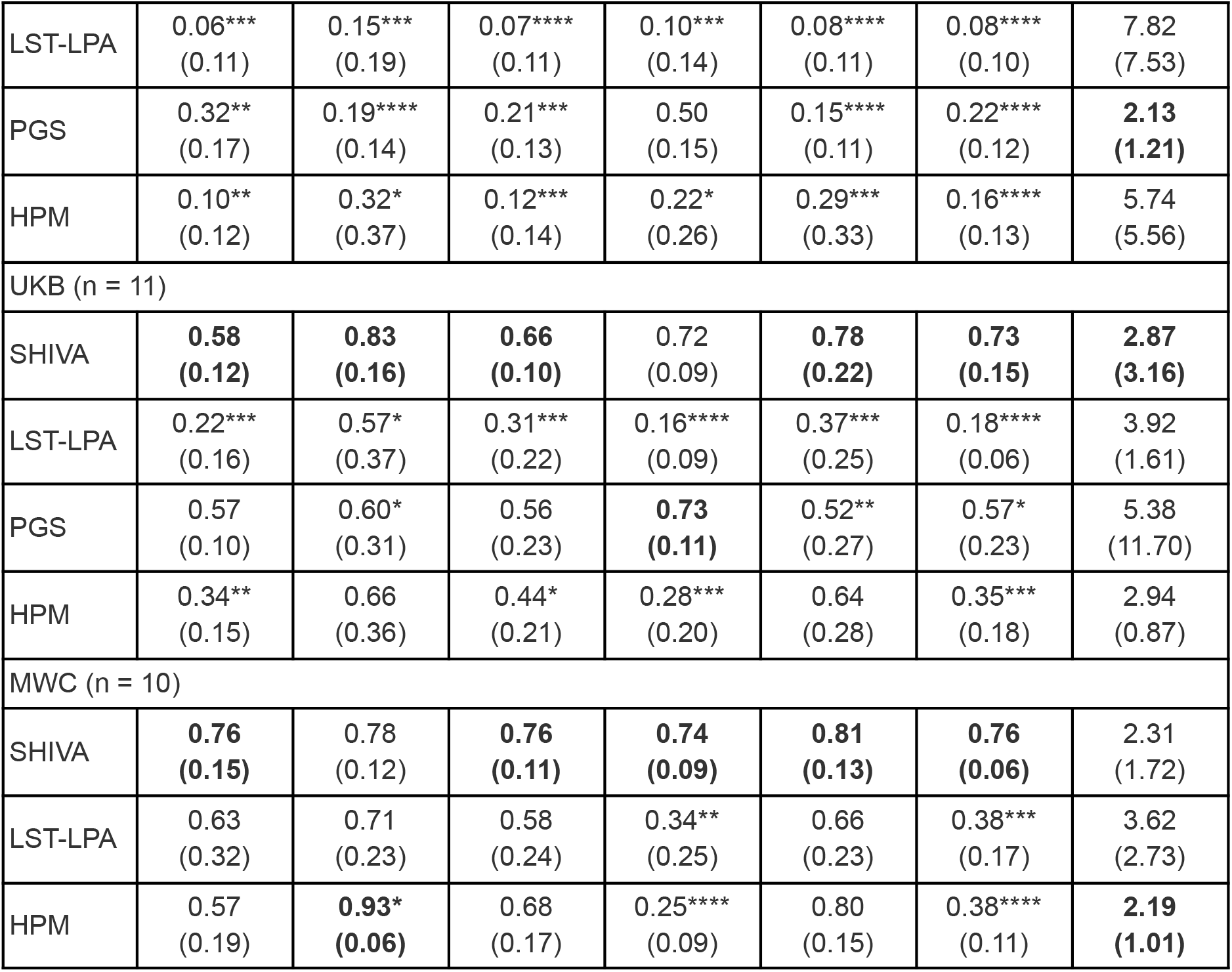
Summary of performance metric comparisons between SHIVA-WMH against the three reference methods separately for each test cohort. Mean and standard deviations (SD) of each metric in each test cohort are shown for SHIVA-WMH and the three reference methods (LST-LPA, PGS, HPM). For each metric in each cohort, best scores are indicated in bold. Asterisk indicates the degree of statistical significance for each paired-t test comparing SHIVA-WMH against each of the reference methods: **** *p* < 0.0001, *** 0.0001 ⩽ *p* <0.001, ** 0.001 ⩽ *p* <0.01, * 0.01 ⩽ *p* <0.05.

They indicate the clear advantage of the SHIVA-WMH detector in the MRi-Share test dataset, both at the voxel- and lesion cluster-level: Our tool shows both higher TPR and PPV than any of the three reference methods, resulting in the significantly higher VL- and CL-Dice scores (all paired t-tests *p* < 0.001 for VL-Dice and *p* < 0.0001 for CL-Dice; Figure 4 and Table 3). FLAIR-only version shows a similar pattern, albeit with slightly lower VL- and CL-Dice scores compared to the multi-modal version and weaker significance when compared against other reference methods (Supplemental Figure 2).

In the MWC test dataset, both multi-modal and FLAIR-only versions of SHIVA-WMH have significantly higher CL-Dice (all *p* < 0.001) and numerically better VL-Dice scores than the LST-LPA or HPM, suggesting that our tool is detecting small lesion clusters better than these tools. While HPM had significantly better VL-PPV than SHIVA-WMH, it came at the cost of lower VL-TPR. In this cohort, we did not perform direct comparison with PGS, since the model had been trained with the subjects we used as the evaluation test set in the present work.

Finally, in the unseen cohort of UKB, with the intermediate level of the overall lesion severity compared to the other two cohorts, multi-modal SHIVA-WMH is clearly superior to the LST-LPA and HPM, with significantly better VL- and CL-Dice scores (both *p* < 0.05, 0.001 for VL- and CL-Dice, respectively), and shows a significantly better CL-Dice than the PGS (*p* < 0.05), in particular showing better VL- and CL-PPV (Figure 5 and Table 3; also see Figure 6 for an example of false positive in PGS). FLAIR-only version has a similar CL-Dice score as the multi-modal version, but shows a slightly worse VL-Dice score in this cohort (Supplemental Figure 2).

## 4. Discussion

Long dismissed as the “normal” radiological finding in the aging brain, WMH is now firmly established as the most common marker of covert cSVD (Debette and Markus, 2010; Wardlaw et al., 2015). In the present work, we leveraged the high-quality research scans from the MRi-Share study (Tsuchida et al., 2021) to demonstrate that young adults in their twenties already exhibit signs of compromised white matter integrity as small punctate WMH. Comparisons of DWI-based metrics in NAWM and manually-traced WMH revealed decreases in both FA and NDI and an elevated MD in these small punctate WMH relative to NAWM. We further described the SHIVA-WMH detector, trained with both the MRi-Share and publicly available MWC dataset from older subjects (Kuijf et al., 2019), with a specific aim to optimize WMH detection across a wider range of WMH burdens than existing methods. We demonstrated the superior performance of our tool relative to three reference methods in accurately segmenting WMH in the test dataset composed of three databases representing young, middle-aged, and older individuals with varying degrees of WMH burden. When performance was compared using Dice score, both at the voxel-by-voxel-level (VL-Dice) and at the level of each lesion cluster (CL-Dice), to evaluate the similarity between the manually-traced and predicted WMH segmentations, our tool had the highest average scores across the test dataset, with the overall VL-Dice and CL-Dice scores of 0.66 and 0.71, respectively. Detailed evaluation in each cohort separately indicated that the SHIVA-WMH detector achieved significantly higher VL- and CL-Dice scores than all three reference methods in the MRi-Share test set, and significantly or nominally higher Dice scores in both UKB and WMC test datasets. It can use either multi-modal inputs of coregistered T1w and FLAIR images or FLAIR image only, although some performance metrics drop slightly for FLAIR-only version. To our knowledge, this is the first automated WMH detection tool that incorporated 3D FLAIR data from young, neurologically asymptomatic adults to train and validate the method in subjects with very mild WMH burden (< 2 mL).

Prior imaging studies have shown that there are significant increases in MD and decreases in FA in WMH compared to NAWM in community-dwelling elderly (Muñoz Maniega et al., 2015; Riphagen et al., 2018; Wardlaw et al., 2015). While few studies have characterized WMH in younger adults, one study that examined the diffusion properties of relatively mild WMH (total lesion volume < 6 mL) in 3D FLAIR scans of neurologically asymptomatic subjects aged between 21 and 60 also reported similar changes, with a significant MD increase and non-significant FA decrease in WMH compared to NAWM (Keřkovský et al., 2019). We extend these earlier observations by demonstrating that the same changes in DTI metrics can already be detected in even milder WMH in young university students. Furthermore, we showed that NDI derived from the NODDI model is sensitive to the microstructural changes in the WMH found in these young subjects, potentially giving more specific insight into the early pathophysiology than DTI metrics alone. It suggests that increased water diffusivity and decreased directionality of diffusion (indicated by MD and FA, respectively) may be driven at least partially by lower myelination or axon density, as indicated by NDI. Although NDI is known to be lower in WMH found in patients with MS (Alotaibi et al., 2021; Mustafi et al., 2019), to our knowledge, this is the first to demonstrate the sensitivity of NDI to WMH in asymptomatic young subjects. Our finding is consistent with early neuropathological work on punctate WMH indicating the reduced myelin content with neuropil atrophy in perivascular tissues in the deep white matter (Fazekas et al., 1998), and indicates that subtle microstructural changes are already detectable with MRI even in young subjects with very low overall WMH load.

Although the present work does not address the functional significance of the small amount of WMH in these young subjects, recent work has highlighted the associations between higher WMH and poorer executive task performance even in young adults aged between 20 and 40 who exhibited a similar degree of WMH burden as MRi-Share subjects in our study (Garnier-Crussard et al., 2020). Further, there is evidence that the amount of WMH found in young adults is associated with several modifiable cardiovascular risk factors, such as body mass index, physical activity, smoking, and alcohol consumption (Williamson et al., 2018). Together, it underscores the importance of accurately charting the early emergence and progression of WMH for a better understanding of its pathophysiology, its genetic and environmental determinants, and ultimately for early intervention.

To this end, we combined the WMH labels of the MRi-Share with the publicly available MWC dataset to develop the SHIVA-WMH detector, based on our prior work that used the 3D Unet-based architecture to detect PVS, another marker of covert cSVD (Boutinaud et al., 2021). Even though there has been an increasing number of studies applying Unet-based models for WMH detection, it has not been used to push the limit of early detection in young, neurologically asymptomatic cohorts. Further, with few exceptions (Tran et al., 2022; Umapathy et al., 2021), most work on automatic segmentation methods for age-related WMH so far has focused on developing and evaluating their tools on more conventional 2D FLAIR images with thick slices (typically 3 to 5 mm). However, there is an inherent limitation in accurately quantifying the amount of small WMH with 2D FLAIR acquisitions, since small lesions out of the plane of acquisition cannot be detected. More modern 3D FLAIR acquisitions with isotropic resolution are known to achieve a better signal/contrast-to-noise ratio and allow greater sensitivity to WMH lesions than 2D acquisitions (Bink et al., 2006; Chagla et al., 2008). They are also increasingly available in large neuroimaging databases in population-based studies (e.g. UKB; Alfaro-Almagro et al., 2018; ADNI-3; Gunter et al., 2017; Rhineland Study; Lohner et al., 2022). Yet, several recent works on age-related WMH detection methods that explicitly evaluated their methods in participants with mild WMH burden (< 5 mL) had not included 3D FLAIR datasets for training or evaluation (Khademi et al., 2021; Mojiri Forooshani et al., 2022; Ong et al., 2022; Rachmadi et al., 2018). Our work is unique in taking advantage of the 1mm isotropic 3D FLAIR scans from young adults and the detailed delineation of every visible WMH in every slice to train the 3D Unet-based model, with the objective to apply our tool in other cohorts with similar acquisitions to accurately describe WMH burden across the adult lifespan.

We evaluated the performance of the SHIVA-WMH detector in the 10 subjects each from the MRi-Share and MWC datasets that had been set aside (i.e. not seen during training), as well as in the 11 additional subjects from the UKB dataset, representing adults from the general population in the age range and with the level of WMH burden in between MRi-Share and MWC. As the reference tools for comparison, we selected one conventional signal intensity-based method (LST-LPA) and two existing Unet-based methods (PGS and HPM) that could be used out-of-the-box (i.e. with pretrained models available for Unet-based methods). The LST-LPA is based on the logistic regression model that uses intensity and location information of FLAIR images to classify WMH. It was originally developed to segment WMH in MS patients (Schmidt, 2017a), but has been applied widely to quantify age-related WMH as well (Ribaldi et al., 2021) and has also been demonstrated to show robust performance across diverse datasets with age-related WMH (Heinen et al., 2019; Vanderbecq et al., 2020). It is also by far the most common algorithm used as a reference method in studies proposing new WMH detection methods (29 out of 37 studies reviewed in Balakrishnan et al. 2021). The PGS is a 2D Unet-based method and the current winner of the MWC 2017 challenge (https://wmh.isi.uu.nl/results/), with the VL- and CL-Dice scores of 0.81 and 0.79 respectively in the held-out test dataset of the challenge (Park et al., 2021). It combines the Unet architecture with what the authors call a multi-scale highlighting foregrounds approach, in which the ground truth labels are down-sampled at multiple scales to allow loss minimization at each layer of the Unet. This approach has the effect of emphasizing the contributions of small lesions and the voxels in the lesion boundaries during the network training, and as a result should improve accurate detection of WMH voxels with high uncertainty due to partial volume effect, including small lesions. Another recently published Unet-based model, HPM uses 3D input like our SHIVA-WMH detector. It was chosen based on the high reported performance in cases of very mild WMH burden in a multi-cohort dataset of 50 subjects with mean WMH volume of ~2 mL, with VL- and CL-Dice scores of 0.84 and 0.72, respectively (Mojiri Forooshani et al., 2022). These scores are the highest among several recent works that explicitly evaluated their methods in participants with mild WMH burden of less than 5 mL (Khademi et al., 2021; Mojiri Forooshani et al., 2022; Ong et al., 2022; Rachmadi et al., 2018). For the purpose of comparison with our tool, it also had the advantage of being trained with multisite imaging datasets that did not include the MWC dataset, allowing for a fair comparison of performance with our tool in this cohort, unlike the PGS, whose training data included MWC testing data set aside in the present study.

Of the three reference methods, LST-LPA had the lowest overall Dice scores, and performed progressively worse in the cohorts with lower overall WMH burden. It had comparable Dice scores as SHIVA-WMH in the MWC test set, but only at the voxel-level. Lesion-wise CL-Dice was significantly worse than SHIVA-WMH even in this test cohort with the largest overall WMH burden. It suggests that while it is able to detect relatively large WMH in subjects with moderate-to high-lesion burden, small lesions found in these subjects are missed by LST-LPA. This observation is consistent with other studies demonstrating the superior performance of Unet-based methods over LST-LPA primarily in subjects with lower overall lesion burden (Khademi et al., 2021; Li et al., 2022). Among the three reference methods, PGS had the highest sensitivity in the MRi-Share and UKB test datasets, with comparable VL- and CL-TPR in UKB and lesion-wise CL-TPR in MRi-Share as SHIVA-WMH, attesting to the stated advantage of their approach. However, the relatively high sensitivity came at the cost of low precision in both test datasets, resulting in the significantly lower VL- and CL-Dice scores in MRi-Share and CL-Dice score in UKB compared to SHIVA-WMH. The lower precision likely results from the 2D input they use for their Unet model, since islands of cortical ribbons on some axial slices are difficult to distinguish from WMH without the 3D context. The underperformance of HPM was somewhat surprising, given their high reported performance in subjects with mild WMH burden and the fact that it has been trained with a large and diverse dataset representing 432 individuals from 4 multicentre studies, using the 3D input for their Unet model as in SHIVA-WMH. Although speculative, we suspect that the reason may be the nature of the training dataset they used: All their training and testing data were from 2D FLAIR with 3 mm slice thickness, which may have limited the advantage of full 3D model. Further, in order to prepare the large number of reference WMH labels to train their model, they used a semi-automated pipeline that generated intensity-based segmentations. Although these labels were then reviewed and manually edited by trained human annotators, it is generally more difficult and time consuming to add lesions missed by the automated method than rejecting false positives during such manual editing, which can result in more conservative labels missing small lesions not detected by the semi-automated method. In contrast, we used the high-quality, high-resolution manual labels for the MRi-Share training dataset consisting of 40 subjects to train a model specific to this cohort, then used the predictions generated by the model in unannotated MRi-Share subjects to iteratively train our model until optimal performance was reached in the manually-traced MRi-Share and MWC validation sets. Such an approach has been recently suggested as one of the effective solutions for enhancing limited high-quality annotations for training DL-based models (Tajbakhsh et al., 2020), here applied specifically to enhance performance for more difficult cases of subjects with very low WMH burden.

The superior performance of the SHIVA-WMH detector relative to other reference methods in the UKB test set is worth being emphasized, as this test set represents data coming from an unseen cohort, and thus constitutes an important test case for the transferability of our detector to cohorts not seen during the training. Further, the UKB dataset has been acquired with a modern scanner with high-quality 3D FLAIR acquisitions similar to MRi-Share, and represents one of the intended target populations to apply our tool in future studies to characterize the full extent of WMH in neurologically asymptomatic adults. Relative to SHIVA-WMH, only PGS showed comparable sensitivity to WMH found in UKB, but at the cost of lower precision, resulting in the lower and more variable Dice scores in this cohort than our tool. It suggests that our tool can predict variable burden of WMH in this cohort more accurately and consistently than the reference tools tested here.

It should be noted that we compared the performance of SHIVA-WMH against PGS and HPM without re-training the latter two methods with the same training data used by SHIVA-WMH, since our aim was to evaluate the direct applicability of these publicly available tools, rather than to test the ability of different Unet-based models to learn new dataset. Although beyond the scope of this study, it is possible that innovations in the model architectures, such as the multi-scale highlighting forgrounds approach in PGS, can further improve small lesion detection when used in combination with the high resolution training data we used. Also, the present work focused on WMH detection in asymptomatic or presymptomatic adults, with the assumption that WMH found in these adults are primarily early stages of age-related WMH of presumed vascular origin, thus combining MRi-Share and MWC dataset for training our model. However, in reality, WMH found in MRi-Share may represent mixed pathology, with lesions in some subjects caused by pre- or sub-clinical inflammatory conditions (Hosseiny et al., 2020). While the model learning the WMH in a given training data is agnostic about their etiology, it can learn any global or local spatial and intensity features of the lesions present in the training dataset. To the extent that there are etiology-specific patterns in WMH appearance and spatial distributions, it is possible that SHIVA-WMH has learnt predominant WMH patterns in the specific training dataset we used, and may be less sensitive to lesions found in other conditions we did not explicitly focus on, such as MS (training dataset for SHIVA-WMH did not contain any incidental MS subjects). Future work should investigate, and if necessary improve, the robustness of our detector across different etiology.

### 4.1. Conclusion

To summarize, we presented the SHIVA-WMH detector, a 3D-Unet based model trained with both MRi-Share and MWC dataset with the specific aim to improve detection of small WMH in asymptomatic or pre-symptomatic adults in population-based studies. Our tool outperformed both a classic WMH segmentation tool (LST-LPA) and existing state-of-the-art Unet-based tools (PGS and HPM) in segmenting small WMH in non-clinical, community-dwelling adults represented by MRi-Share and UKB. Our tool can effectively segment WMH across a wider range of WMH burden than existing methods, and thus can be a valuable tool for studies aiming to characterize the emergence and progression of WMH lesions. Such studies are essential for understanding the pathophysiology and early-life factors associated with the most common etiology of WMH in the population, namely cSVD. Our demonstration of altered diffusion properties of small WMH in MRi-Share, bearing the hallmark of compromised microstructural integrity similar to those found in WMH of older subjects or MS patients, also underscores the importance of early detection and intervention. To encourage more research on the early detection and characterization of WMH, we make the SHIVA-WMH detector freely and openly available at (https://github.com/pboutinaud/SHIVA_WMH).

## Data/code availability statement

MRi-Share data used in this study cannot be shared through a public repository due to French regulations regarding sharing of the medical imaging data. However, de-identified data can be requested to the i-Share Scientific Collaborations Coordinator with a letter of intent (explaining the rationale and objectives of the research proposal), and a brief summary of the planned means and options for funding. MICCAI 2017 WMH segmentation challenge dataset used in the present work is freely and publicly available at the challenge homepage (https://wmh.isi.uu.nl/data/). This work also used the neuroimaging dataset obtained from the UK Biobank Resource (application number 18359 and 94113).

Source codes for the statistical analysis presented in the manuscript are available on GitHub (https://github.com/atsuch/SHIVA-WMHpaper). SHIVA-WMH detector presented in this work is also publicly available at https://github.com/pboutinaud/SHIVA_WMH.

## CRediT authorship contribution statement

**Ami Tsuchida:** Conceptualization, Formal analysis, Investigation, Data Curation, Writing-Original Draft, Visualization; **Philippe Boutinaud:** Conceptualization, Methodology, Software, Writing-Review & Editing; **Violaine Verrecchia**: Data Curation; **Christophe Tzourio:** Funding acquisition, Writing-Review & Editing; **Stéphanie Debette:** Funding acquisition, Project administration, Writing-Review & Editing; **Marc Joliot:** Conceptualization, Supervision, Project administration, Writing-Review & Editing.

## Acknowledgements

We would like to acknowledge Prof. Bernard Mazoyer for the initial conception of the project to characterize early signs of white matter anomalies in the young subjects of MRi-Share and his reviewing of raw T1w and FLAIR images in this database to select 50 subjects with ranging amount of both WMH and PVS. We are also indebted to the following individuals for their invaluable contribution to the MRi-Share project: Serge Anandra, Amandine André, Gregory Beaudet, Christophe Bernard, Bruno Brochet, Aurore Capelli, Claire Cardona, Arnaud Chaussé, Christophe Delalande, Vincent Durand, Louise Knafo, Morgane Lachaize, Alexandre Laurent, Hugues Loiseau, Elena Milesi, Marie Mougin, Maylis Melin, Guy Perchey, Clothilde Pollet, Thomas Tourdias, Cécile Marchal, Guillaume Penchet, Cécile Dulau, Igor Sibon, Sabrina Debruxelle, Sophie Auriacombe, Caroline Roussillon, Nicolas Vinuesa, and the i-Share “relay” students. We would also like to express our gratitude to Paul Matthews (Imperial College, London, UK) and to the personnel of the UK-Biobank imaging center at Stockport (UK) for their help while designing the MRi-Share image acquisition protocol, and to Maxime Descoteaux (Sherbrooke University, Canada) for his help in implementing the DWI processing and QC pipelines. Finally, the authors would like to express their gratitude to the 1,870 students of the Bordeaux University who gave their consent to participate in MRi-Share.

