## Supplemental Material for "Early detection of white matter hyperintensities using SHIVA-WMH detector"

### Enhancement of SHIVA-WMH with training data derived from a model trained with MRi-Share (‘MRi-Share-specific’ model)

Our first “base” model (dark green in Supplemental Figure 1) was trained with all the available training datasets that included 40 MRi-Share and 50 MICCAI2017 WMH challenge (MWC) data with the manual tracing of WMH. We suspected the imbalance in the amount of WMH found in the two cohorts may bias the model toward learning better the large WMH found in MWC. To gauge how well the model can learn to segment milder forms of WMH in MRi-Share data when focusing the training on this cohort alone, we trained the same model with 40 MRi-Share data only (“MRi-Share-specific” model, yellow in Supplemental Figure 1). We compared the performance of this “MRi-Share-specific” model and the “base” model in MRi-Share by comparing the average of voxel-level (VL-) and cluster-level (CL-) Dice scores for each subject in the validation set of each training fold. The VL- and CL-Dice scores were averaged to evaluate the performance to maximize segmentation accuracy at both individual voxels and lesion clusters. As shown in Supplemental Figure 1, the “MRi-Share-specific” model could achieve higher average VL- and CL- Dice scores in MRi-Share than the “base” model despite the smaller overall training data size, consistent with the bias introduced when combining the MRi-Share and MWC training datasets. Unsurprisingly, the performance of the “MRi-Share-specific” model was poor in the MWC datasets, indicating that learning WMH from MRi-Share data only does not transfer well to more severe forms of lesions found in MWC.

In order to improve the model performance in subjects with only mild lesions without compromising its performance in subjects with higher lesion load, we generated more labels of WMH in a random batch of 100 MRi-Share subjects without manual WMH tracing using the “MRi-Share-specific” model. We filtered out the top and bottom 5 subjects based on the predicted WMH load to remove the influence of outliers and used the predicted WMH labels for the remaining 90 subjects to enhance our original training dataset (40 MRi-Share + 50 MWC). The model with the enhanced training data (“enh90”) was trained using the “base” model as initial weights to speed up the learning. This “enh90” model brought the average VL- and CL-Dice scores in MRi-Share comparable to those of “MRi-Share-specific” model, and even slightly improved the performance in MWC relative to the “base” model. Encouraged by the results, we repeated the procedure with the prediction generated in a new batch of 270, 300, and 400 MRi-Share subjects successively (“enh270”, “enh300”, and “enh400”, respectively), each time using the previous model to initialize the weights. Because there were no noticeable improvements after the second iteration of the enhancement, we selected the “enh270” model and used it as the SHIVA-WMH detector in the main text.

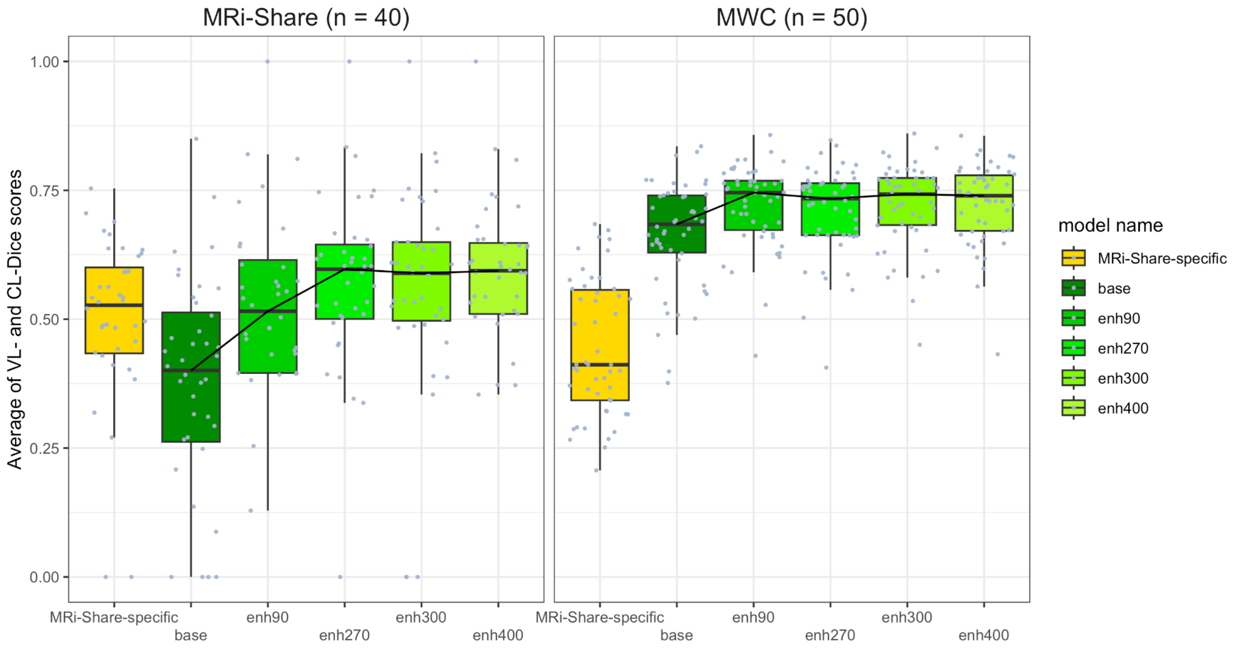


**Supplemental Figure 1. Averaged voxel- and cluster-level Dice scores of the "MRi-Share-specific" model compared to the "base" model and successive models with enhanced training datasets ("enh90", "enh270", "enh300", "enh400") in the validation set in each training fold, collapsed across the folds.**
*Left Panel*: Performance comparison in the 40 MRi-Share training dataset. *Right Panel*: Performance comparison in the 50 MWC training dataset, but note that the 'MRi-Share-specific' model did not include these subjects in its training. Median values of the 'base' model and successive models with enhanced training datasets are connected to facilitate the comparison of improvements with each iteration of enhancements.

### Development of FLAIR-only version and performance comparison

The FLAIR-only version of SHIVA-WMH was trained with the same original 90 training dataset (40 MRi-Share and 50 MWC) enhanced with 270 additional WMH labels generated from the "MRi-Share-specific" model that used the T1w and FLAIR images in unannotated MRi-Share subjects to segment WMH. Instead of being trained iteratively like in the multi-modal version, however, this FLAIR-only version was trained from scratch, using the Glorot initializer for the initial weights.

We evaluated its performance relative to the three reference methods in an identical manner as the primary multi-modal version described in the main text, by performing a series of paired t-tests comparing each reference method against the FLAIR-only version. Supplemental Table 1 summarizes the performance metrics of the FLAIR-only version across the 31 test-set subjects. Supplemental Figure 2 shows the cohort-specific comparisons of VL- and CL-Dice scores against the reference methods. Together, they indicate that the FLAIR-only version has similar advantages over other methods as the primary multi-modal version, although some metrics are slightly lower overall.

**Supplemental Table 1. Comparison of SHIVA-WMH (FLAIR only) against three reference methods across the 31 test subjects for each performance metric.**

The mean and standard deviations (SD) of each metric across all the test subjects are shown for the FLAIR-only version of SHIVA-WMH and the three reference methods (LST-LPA, PGS, HPM). For each metric, the best scores are indicated in bold. Asterisk indicates the degree of statistical significance for each paired-t test comparing SHIVA-WMH against each reference method: **** $p$ < 0.0001, *** 0.0001 ⩽ $p$ <0.001, ** 0.001 ⩽ $p$ <0.01, * 0.01 ⩽ $p$ <0.05.

| | Mean (SD) | | | | | | |
|---|---|---|---|---|---|---|---|
| | VL-TPR | VL-PPV | VL-Dice | CL-TPR | CL-PPV | CL-Dice | HD95 |
| All (N = 31[a]) | | | | | | | |
| SHIVA (FLAIR only) | **0.55 (0.22)** | **0.76 (0.22)** | **0.59 (0.18)** | **0.60 (0.16)** | **0.88 (0.14)** | **0.71 (0.14)** | **3.15 (3.18)** |
| LST-LPA | 0.30*** (0.32) | 0.48** (0.36) | 0.32*** (0.29) | 0.20**** (0.20) | 0.37**** (0.31) | 0.21**** (0.17) | 4.55* (3.93) |
| PGS | 0.45 (0.18) | 0.41**** (0.32) | 0.39 (0.26) | 0.62* (0.17) | 0.34**** (0.28) | 0.40**** (0.26) | 3.83 (8.48) |
| HPM | 0.34** (0.24) | 0.64 (0.38) | 0.42* (0.29) | 0.25**** (0.19) | 0.58*** (0.33) | 0.30**** (0.17) | 3.45 (3.24) |

[a] Comparison with PGS was performed in 21 test subjects that excluded subjects from MWC.

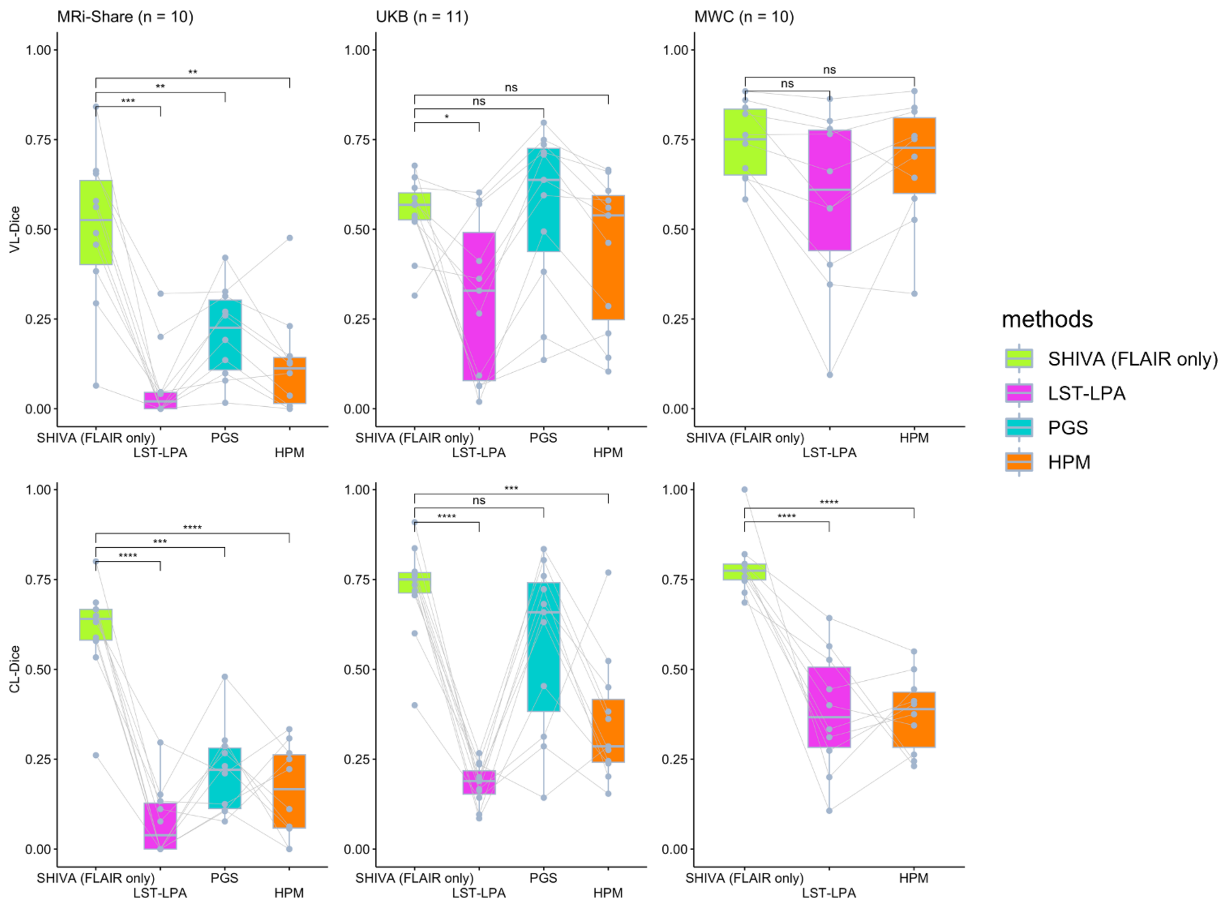


**Supplemental Figure 2. Voxel- and Cluster-level Dice scores of SHIVA-WMH (FLAIR only) compared with LST-LPA, PGS, and HPM tools in the test-set subjects in each cohort.** Comparisons of VL-Dice (*top row*) and CL-Dice (*bottom row*) scores between the FLAIR-only version of SHIVA-WMH against the reference tools (LST-LPA, PGS, and HPM) are shown, separately for MRi-Share (n = 10), UKB (n = 11), and MWC (n = 10) test subjects. Asterisk indicates the degree of statistical significance for each paired-t test comparing SHIVA-WMH against each reference method: **** $p < 0.0001$, *** $0.0001 \leqslant p < 0.001$, ** $0.001 \leqslant p < 0.01$, * $0.01 \leqslant p < 0.05$, ns $p \geqslant 0.05$.
